# Submicron-Sampling of Living Cells by Macrophages

**DOI:** 10.1101/2025.04.10.648051

**Authors:** Amy C. Fan, Rukman Thota, Nina Serwas, Vivasvan S. Vykunta, Kyle Marchuk, Megan K. Ruhland, Lauren Liu, Grace Johnson, Austin Edwards, Matthew F. Krummel

## Abstract

An effective immune system must sample and appreciate healthy-self identity to prevent autoimmunity and to contrast to pathogenic insults^1–3^. Self-proteins are presented to T cells in the thymus during immune cell development^2,3^, and must be presented throughout the body to both maintain regulatory T cell populations^4–6^ and provide a tonic signal to maintain conventional T cells over time^7–9^. The ready observations of continuous apoptosis in some organs together with the ingestion of that material by myeloid populations has led to a conventional understanding of ongoing cell-death as a major source of self-antigens^10^, complemented in some situations by uptake of free-floating cell-derived vesicles. Here, we used a series of companion imaging and vesicular labeling technologies to reveal an alternate process undertaken by macrophages that results in non-destructive and direct sampling of living cells. The process requires cell-cell contact, does not require caspase activation, and takes place via a trogocytosis-like stretching of the target cell into the macrophage, leading to the generation of submicron-sized vesicles containing cytoplasm. Using a high-dimensional flow-based method for labeling vesicles ingested under this versus other conditions, we find that live-sampled material is distinctly processed, is poorly subject to fusion with lysosomes, and produces ensuing differential effects on the presentation of those to CD4 versus CD8 T cells. Disrupting this trafficking by redirecting antigen to the lysosome significantly reduced the associated macrophage-mediated priming of CD8 T cells. This demonstrates an important and substantial sampling of living cells by the immune system, with clear consequences for maintaining the border of immunity.

## Tissue and tumor-associated proteins are routinely sampled by myeloid cells

Antigen presenting cells (APCs) must continuously survey tissues by ingesting and processing antigens for presentation to direct antigen-specific T cell responses^11^. In the course of studying a selection of tissue-specific sites of tolerance, including in tumors, we expressed the fluorescent protein ZsGreen in multiple non-inflammatory settings across a range of tissues with slow turnover rates (Scbg1a1-Cre-ERT2; airway epithelium)^12^ or faster turnover (K14-Cre skin, tumors, Villin-Cre intestinal epithelium). ZsGreen is both bright and very stable, permitting superior long-lived tracking^1,10,24^. Cytosolic ZsGreen expressed under these promoters all routinely illuminated substantial CD45^+^ immune populations that contained numerous submicron-sized vesicular puncta of ZsGreen (**Fig. 1a, b)**, consistent with ingestion of tissue-associated protein. When CD45^+^ cells bearing ZsGreen^+^ were isolated from these healthy tissues, a variety of myeloid cells were highlighted (**Fig. 1c, d; Extended Data Fig. 1a,b; Supplementary Fig. 2**), including dendritic cells and neutrophils but with macrophages being the most consistently loaded. These loading frequencies also mirror previous characterization of myeloid sampling of skin^13^ and tumor cytoplasts^14^.

**Figure 1:**
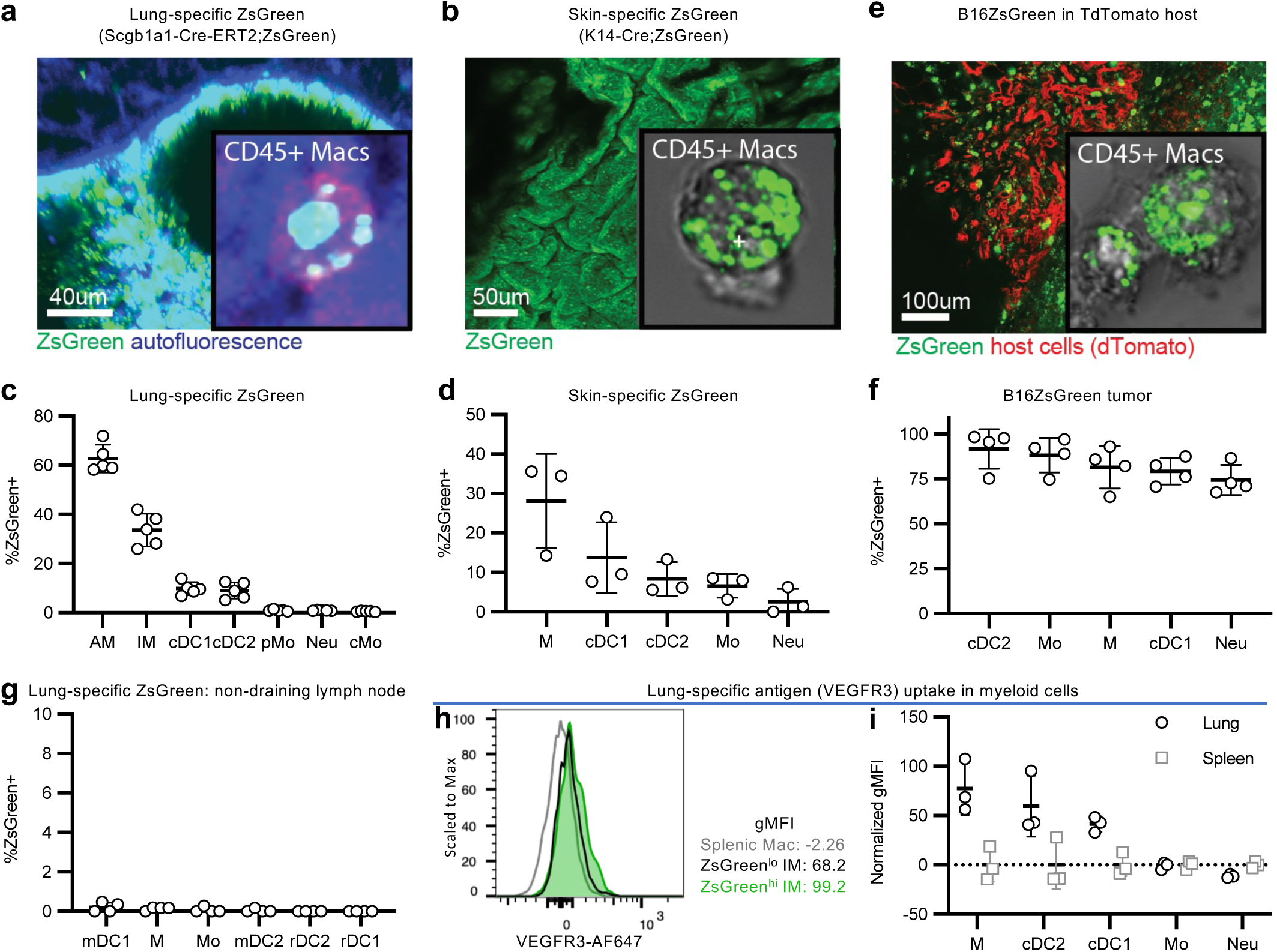
Myeloid cells sample proteins from healthy and tumor tissues. **a-b**, Lungs (**a**) or skin (**b**) taken from mice expressing ZsGreen under the Scgb1a1 (**a**) or K14 (**b**) Cre with insets of in situ macrophages labeled with CD45 (**a**) or sorted CD45+ Macrophages (**b**) showing ZsGreen puncta inside macrophages. **c-d,** Isolation of lung myeloid cells from Scgb1a1-Cre-ERT2;ZsGreen mice (**c**) or skin myeloid cells from K14-Cre;ZsGreen mice (**d**) shows specific uptake in tissues prominently in macrophages. Lymphocytes were routinely ZsGreen negative. **e,** B16 melanoma cells expressing ZsGreen transplanted into mice ubiquitously expressing TdTomato with inset of CD45+ macrophages. **f**, Isolation of tumor-associated myeloid cells shows broad uptake of ZsGreen across myeloid cell subsets. **g**, Isolation of inguinal LN myeloid cells from Scgb1a1-Cre-ERT2;ZsGreen mice shows no uptake in distant lymph nodes. **h,** Intracellular staining for airway specific VEGFR3 protein in lung (black) compared to splenic (gray) myeloid cell populations with ZsGreen^hi^ lung macrophages (green) further enriching for VEGFR3 signal. **i,** VEGFR3 MFI in lung and splenic myeloid cell populations normalized to average MFI of splenic myeloid cells. Representative of 3 experiments, n = 3-6 mice per experiment. AM, alveolar macrophage; IM, interstitial macrophage; pMo, patrolling monocyte; Neu, neutrophil; cMo, conventional monocyte; M, macrophage; Mo, monocyte, mDC2, migratory cDC2; mDC1, migratory cDC1; rDC1, residential cDC1; rDC2, residential cDC2

These observations do not result from misexpression of transgenes in host tissues as demonstrated by the following series of observations. First, we compared the ZsGreen fluorescence intensity of lung myeloid cells in mice expressing ubiquitous ZsGreen (“B-actin-Cre;ZsGreen”), lung-specific ZsGreen (“Scgb1a1-Cre-ERT2;ZsGreen”), or no ZsGreen (C57BL/6 wildtype, “B6 WT”). In each myeloid cell population, ZsGreen fluorescence intensity in the Scgb1a1-Cre-ERT2;ZsGreen mice was greater than in B6 WT mice but less than in B-actin-Cre;ZsGreen mice (**Extended Data Fig. 1b**), consistent with accumulation of exogenous ZsGreen protein as opposed to cell-autonomous ZsGreen expression. Second, normal bone-marrow transplanted into mice expressing ZsGreen gave rise to donor-derived myeloid cell populations that were bright for ZsGreen (**Extended Data Fig. 1c; Supplementary Fig. 3a**). Third, in mice ubiquitously expressing TdTomato transplanted subcutaneously with B16F10 tumors expressing ZsGreen (“B16-ZsGreen”), host CD45^+^ immune cells contained tumor-derived vesicular ZsGreen puncta and a significant proportion of tumor-associated myeloid cells ingested ZsGreen (**Fig. 1e,f, Supplementary Fig. 3b**). Finally, whereas we had previously observed trafficking of tissue-specific ZsGreen to tissue draining lymph nodes^13^, ZsGreen was not detected in the non-draining lymph node of Scgb1a1-Cre-ERT2;ZsGreen mice (**Fig. 1g**), indicating tissue-specific uptake.

For tumor antigens, it was previously found that tumor antigens were co-packaged in macrophage vesicles, together with tumor-derived ZsGreen tracer^13^. To determine if other non-tumor self-antigens in healthy tissues with low turnover like epithelium were similarly taken up in these myeloid cell populations, we examined the levels of VEGFR3 and PLVAP proteins, which are specifically expressed on the cell surface of non-hematopoietic cells in the lung. When we examined the cellular components from the lungs of Scgb1a1-Cre-ERT2;ZsGreen mice by intracellular flow-cytometry, we found that local myeloid cell populations from the lung, but not distant splenic myeloid cells, contained these self-proteins **(Fig. 1h, i; Extended Data Fig. 1d; Supplementary Fig. 3c)**. Further, ZsGreen^hi^ myeloid cells were brighter for the VEGFR3 and PLVAP stains as compared to ZsGreen^lo^ myeloid cells (**Fig. 1h; Extended Data Fig. 1d**). Therefore, tissue-associated myeloid cells regularly sample tissue-associated self-proteins, both normal and tracer.

## Substantial antigen sampling from live cells by macrophages

Several mechanisms of uptake might contribute to myeloid cell sampling from tissues in vivo, with the most heavily studied to date being phagocytosis of dead cells and endocytosis of exosomes. To formally interrogate how myeloid cells can obtain intracellular material from nearby cells, we established an in vitro assay for sampling using donor cell populations, grown at ∼99% viability, in log phase. We co-cultured bone marrow derived macrophages (BMDMs) with those ZsGreen-expressing target, especially focusing on two model target cells: B16-ZsGreen melanoma cells which in these conditions minimally produce exosomes, or primary mouse embryonic fibroblasts isolated from mice ubiquitously expressing ZsGreen (“MEF-ZsGreen”). From this co-culture, we detected significant ZsGreen^+^ uptake from both target cells into the BMDM, with the intensity of that ZsGreen being hundreds of times lower than the intensities of intact donor cells, consistent with partial sampling as opposed to complete engulfment (**Fig. 2a, Supplementary Fig. 4a**). When those ZsGreen^+^ BMDMs were sorted for imaging by high-resolution spinning disk confocal microscopy, we readily visualized internalized submicron-sized ZsGreen^+^ puncta **(Fig. 2b-c; Extended Data Fig. 2a)**.

**Figure 2:**
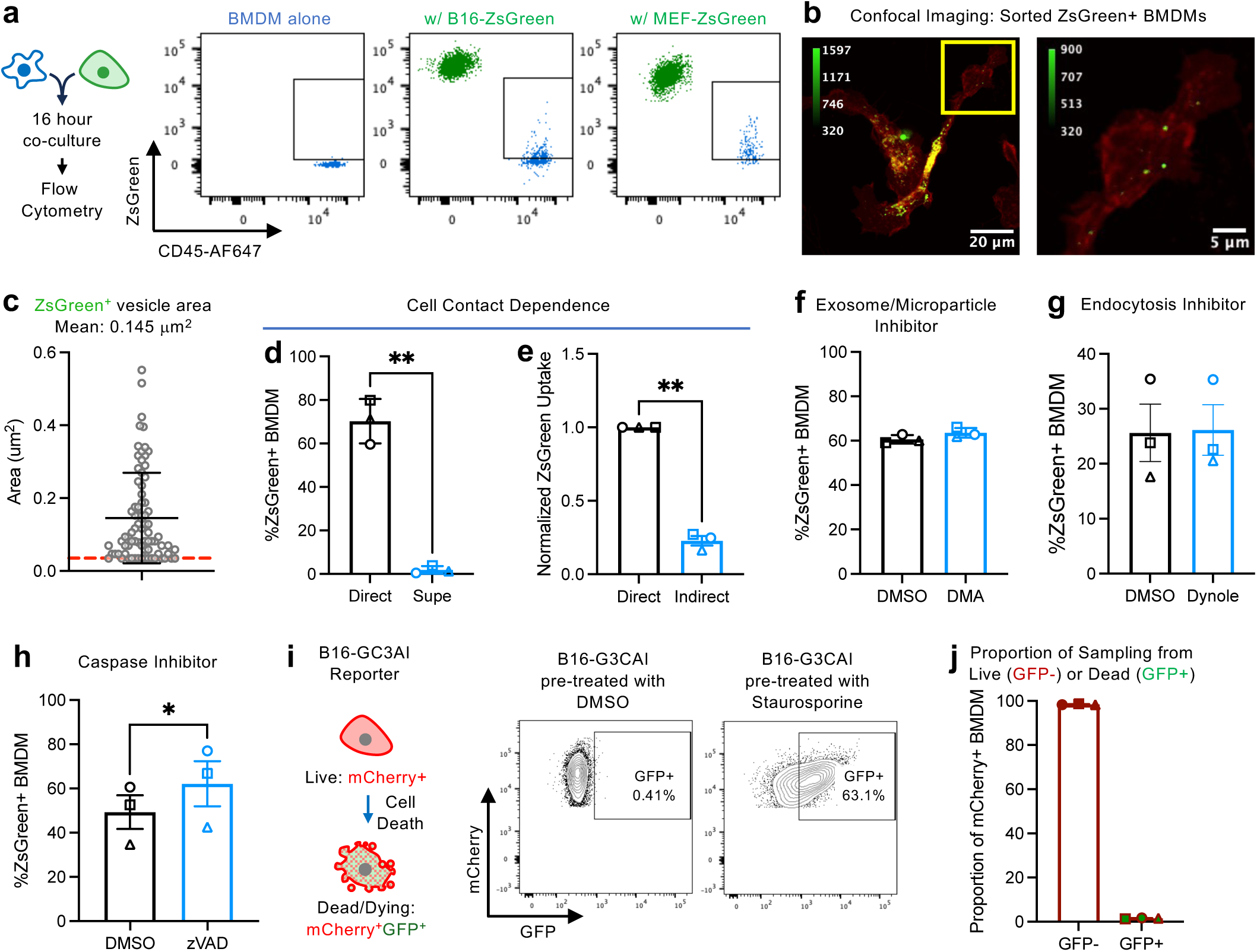
Live cells can be sampled in a cell-contact dependent manner without caspase activation. **a**, BMDMs and ZsGreen-target cells are co-cultured for 16 hours prior to evaluation for ZsGreen uptake by flow cytometry. **b**, Sorted ZsGreen^+^ BMDMs with membrane Tomato showcasing ZsGreen^+^ puncta. (**c**) Area of individual ZsGreen^+^ vesicles. n = 77 vesicles quantified from 23 cells. Shown are mean +/- s.d. **d-e**, BMDMs were co-cultured directly with B16s (“Direct”), with B16 supernatant (**d**, “supe”), or with a transwell insert containing B16s (**e**, “Indirect”)). **f-h**, In vitro co-cultures were treated with indicated inhibitors or DMSO vehicle control. **i,** Induction of GFP in B16-F10 expressing GFP caspase 3 activity indicator (B16-GC3AI) treated with DMSO or Staurosporine. **j**, BMDM co-culture with B16-GC3AI demonstrates uptake is predominantly from live mCherry^+^GFP^-^ target cells. n = 3 biological replicates. Shown are mean of technical replicates +/- s.e.m. Two-sided paired t-test. * p < 0.05, ** p < 0.01. MEF, mouse embryonic fibroblast.

We then evaluated whether antigen sampling from live cells in this setting is mediated through uptake of soluble particles and/or was cell-contact dependent, where the former in particular is expected for the ingestion of free exosomes and perhaps apoptotic blebs. Culture of BMDMs with either B16-ZsGreen derived supernatant containing the limited number of exosomes produced in a 48-hour period or B16-ZsGreen cells that were separated from the BMDM by a transwell both significantly reduced uptake (**Fig. 2d, e; Extended Data 2b**). Additionally, treatment of cultures with inhibitors of either exosome/microparticle release or endocytosis both did not reduce uptake (**Fig. 2f, g; Extended Data 2c-f; Supplementary Fig. 4b, c**). Therefore, although endocytosis of soluble material such as exosomes and microparticles may contribute modestly, this suggested a distinct dominant mechanism of uptake of live cell-associated material in this setting, requiring cell-contact.

Since an existing paradigm suggests that phagocytosis of apoptotic bodies, termed efferocytosis but heretofore just “phagocytosis”, is the major mechanism by which tissues donate material to surveilling APCs^15–17^, we sought to test whether cell death is necessary for the substantial sampling in our system. When we treated the co-culture with the caspase inhibitor zVAD, we found no effect upon ZsGreen uptake, a result which does not support that this uptake relies on cell death (**Fig. 2g, Extended Data Fig. 2f,g; Supplementary Fig. 4d**). To study more directly whether the material ingested into macrophages came from cells undergoing apoptosis, we used a B16-F10 model cell line expressing constitutive mCherry and a split GFP caspase 3 activity indicator which fluoresces only when cleaved, as during apoptosis^18^ (“B16-GC3AI”) (**Fig. 2i**). Use of this confirmed that the B16-GC3AI cells that went into our assays were highly viable (0.12% GFP^+^), whereas apoptotic cell death induced by Staurosporine resulted in GFP reporter fluorescence (>63% GFP, **Fig. 2i**). When these reporter-expressing cells were co-cultured with BMDM, we found that the majority of the mCherry-positive cells (those that had taken up material from the donor cells) lacked GFP expression compared to uptake following apoptosis (1% for live sampling, versus 11% following staurosporine; **Fig. 2j, Extended Data Fig. 2h**). Together, these results further support that an alternative sampling pathway, beyond those involving apoptosis, exists for obtaining material from live cells.

## Live-Imaging of Live-sampling

Given the small size of particles we found within macrophages, we adopted high-resolution methods to directly image the time-course of B16-ZsGreen cells interacting with BMDMs expressing membrane-TdTomato in a co-culture. We variously used lattice light sheet imaging with isotropic resolution of approximately ∼220nm, or the Nikon Spatial Array Confocal (NSPARC), detector system which utilizes an ultra-low noise detector array with approximate lateral and axial resolutions of approximately ∼212nm and ∼424nm, respectively. Importantly, both technologies can sample live full cell volumes over time with reduced phototoxicity.

Using NSPARC, we observed target cell-BMDM interactions that spontaneously result in the stretching of a protrusion of the target cell into the BMDMs. This was frequently followed by the separation of a distinct ZsGreen^+^ vesicle (**Fig. 3b**). In a representative movie (**Fig. 3b, Supplementary Movie 1**), this entire process took <10 minutes from imaging of the initial ZsGreen^+^ protrusion until the time when a protrusion was no longer visible from the target cell, although we registered the actual abscission of the vesicle within a single frame (30 seconds; **Fig. 3a, b**) followed by complete loss of the remaining tether, either into the vesicle or back into the donor cell. We confirmed, at the end of the process, that very small amounts of target cell material were physically inside the BMDM by analyzing the same data in YZ and XZ projections (**Extended Data Fig. 3a**). Consistent with the size ranges we have observed in vivo^13^, these separated vesicles were small, ranging from diffraction limited (<∼0.02μm^3^) to ∼0.05μm^3^ (**Fig. 3c**). In some cases, we also saw a protrusion pulled into the BMDM but then retracted without detectable pinching of a vesicle, which might either have led to extremely tiny ingestion or was simply abortive. Post-ingestion, ZsGreen^+^ puncta moved within the cytoplasm (**Extended Data Fig. 3b**). These data suggest that live-cell cytosolic material can be sampled in a contact-dependent, trogocytosis-like manner.

**Figure 3:**
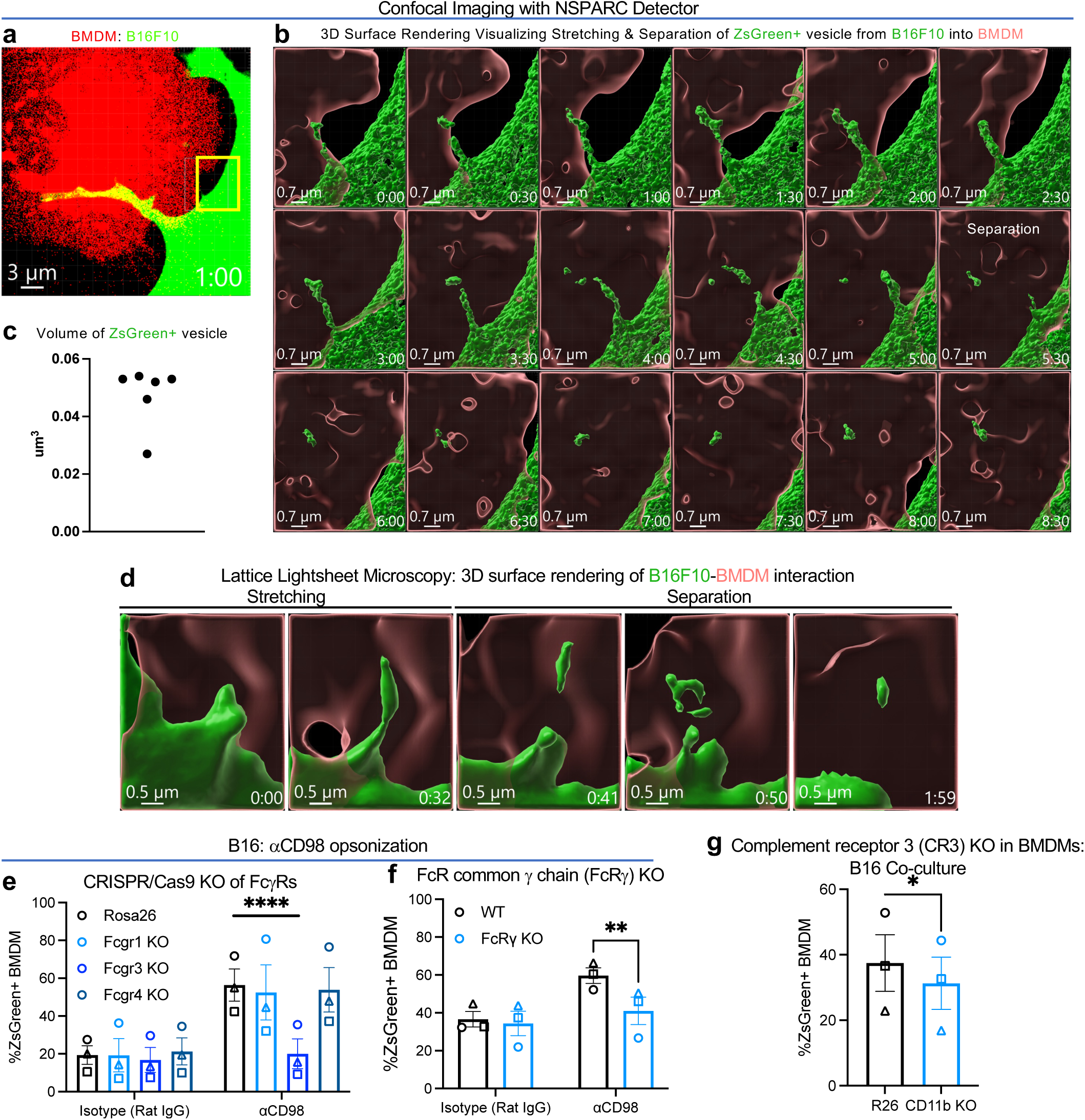
Trogocytosis-like sampling enables macrophages to ingest cytosolic protein from live cells. **a-b**, Confocal live imaging of TdTomato^+^ BMDM and ZsGreen^+^ B16F10 interactions using the NSPARC detector. Time shown as m:ss. **a,** Two-color volume rendering displaying interaction between TdTomato^+^ BMDM and ZsGreen^+^ B16F10. Yellow box highlights the region of interest displayed in **(b). b,** Three-dimensional surface rendering from ROI in **(a)** visualizing the formation and separation of a vesicle from ZsGreen+ B16 into TdTomato+ BMDM. Z-stacks within series were captured in 30 sec intervals. **c,** Volume measurements of separated vesicles from frames 13-18 in **(b)** using Imaris object statistics. **d,** Lattice light-sheet imaging of TdTomato^+^ BMDM and ZsGreen^+^ B16F10 capturing the stretching and separation of ZsGreen^+^ vesicle from target cell. Time shown as m:ss. **e-f,** Antibody-opsonized B16-ZsGreen target cells were co-cultured with either (**e**) BMDMs targeted at the *Rosa26* control locus or receptor loci using CRISPR-Cas9 RNP or (**f**) BMDMs isolated from wild-type B6 mice or FcRγ KO mice. **g**, BMDMs targeted at the *Rosa26* control locus or *Itgam* locus encoding CD11b were co-cultured with B16-ZsGreen target cells. n = 3 biological replicates. Shown are mean +/- s.e.m. Two-way ANOVA with Sidak’s multiple comparison or two-sided paired t-test. * p < 0.05, ** p < 0.01, **** p < 0.0001.

To confirm that this process was not an artefact of the detection method and to take advantage of faster frame-rates, we also leveraged lattice light sheet (LLS) microscopy^19^ with its isotropic high resolution, which again captured the stretching and separation of ZsGreen^+^ vesicles, this time again visible across a single frame (now ∼8 seconds; **Fig. 3d, Extended Data Fig. 3d, Supplementary Movie 2**). Although ingested vesicles were occasionally identified using spinning disk confocal (**Extended Data Fig. 3c**), it is possible that the small size of these vesicles (limiting total fluorescent yield) and the lower sampling rate of this ingestion method limits our ability to capture this biology with older technologies. This also provides one possible explanation why this process has not yet been described or extensively studied by conventional microscopic methods.

## Receptor-mediated mechanisms mediate sampling from live cells

Although multiple receptor-ligand interactions are likely to contribute to this process, we first asked whether previously described mechanisms such as antibody-opsonization or complement receptor 3 (CR3) engagement couple facilitate sampling from live cells. To this end, we found that, pre-incubation with antibodies to surface protein CD98 increased ZsGreen uptake (**Fig. 3f**). Consistent with previously reported binding specificity of Rat IgG1κ to Fcγ receptor 3 (FcγR3, CD16)^20^, this enhancement required FcγR3 binding and signaling (**Fig. 3f-i, Extended Data Fig. 4a-c**). Pre-incubation with antibodies to another abundant surface protein, CD29, similarly increased uptake in an FcγR-dependent manner (**Extended Data Fig. 4d-f**). Moreover, pre-incubation of ZsGreen target cells with normal mouse serum or isolated IgG prior to co-culture with BMDMs increased ZsGreen uptake (**Extended Data Fig. 4g,h**), suggesting that even weakly cross-reactive collections of antibodies amplifies sampling from live cells. Furthermore, we found that disruption of complement receptor CR3 (CD11b) in BMDM decreases the frequency and intensity of ZsGreen uptake in B16 and MEFs (**Fig. 3h, Extended Data Fig. 5a-c**), similar to findings that synaptic pruning by microglia involves these receptors^21^.

Next, to identify other surface proteins beyond CD11b and FcγR involved in this process, we used NicheNet^22^ to predict ligand-receptor interactions between our two model target cells and BMDMs (**Extended Data Fig. 5d**). Because CD11b also has established roles in phagocytosis, we prioritized shared receptors with known phagocytic functions^23^, confirmed their protein, and evaluated their contribution to ZsGreen uptake. Of the eight candidate receptors that we examined, only KO of CD93, a C-type lectin receptor, showed a modest but significant decrease in ZsGreen uptake (**Extended Fig. 5e-g**). These results indicate that multiple surface interactions contribute to uptake, with CD11b and CD93 playing partial roles.

We next examined signaling pathways that link receptor engagement to uptake using small-molecule inhibitors. We targeted signaling downstream of CD11b (Src, Syk, PI3K), small GTPases important for vesicle trafficking (Arf6, Cdc42, Rac), and actin nucleation proteins (Arp2/3, formins). Inhibiting Src signaling (PP1), PI3K signaling (GDC-0941), and Arp2/3 (CK-666) decreased but did not completely abrogate uptake (**Extended Data Fig. 6**). These results suggest that receptor mediated activation of Src and PI3K, leading to Arp2/3-driven branched actin assembly, supports vesicle formation. In contrast, Syk, small GTPases, and formin-driven linear actin may be dispensable or redundant in our system. Altogether, these data indicate that pleiotropic mechanisms contribute to this contact-dependent sampling process that nevertheless relies on local activation cues at the cell-cell interface.

## Tracking sampled material using high-dimensional intracellular vesicle flow cytometry

We sought to determine whether this mechanism has any distinct functional downstream consequences for the vesicle or cargo by first interrogating its fate as compared to material obtained by phagocytosis or endocytosis. To facilitate a high-dimensional analysis of the organelles that derived from different modes of cell sampling, we developed a 10-parameter flow-based organelle profiling method, loosely based on a previous phagoFACS^24^, here capable of immunophenotyping of multiple intracellular compartments (**Fig. 4a**, see **Methods**). Importantly, we incorporated a cell-surface biotinylation step prior to lysis, to allow definitive discrimination of surface-derived membrane, and we also stained the preparation with amine-binding CellTraceViolet to focus analysis upon protein-associated intracellular organelles (**Fig. 4b**, **Supplementary Fig. 5a, see Methods**). Reassuringly, markers for intracellular organelles such as Early Endosome Antigen 1 (EEA1) and RAB7 are exclusively detected in the streptavidin-negative material (**Fig. 4c**). In CellTraceViolet^+^ intracellular vesicles isolated from BMDMs, we were able to distinguish ZsGreen^+^ antigen-containing vesicles from ZsGreen^-^ antigen-free vesicles, and these were exclusively found in BMDMs that had been co-cultured with ZsGreen-expressing target cells (**Fig. 4d, Supplementary Fig. 5b**).

**Figure 4:**
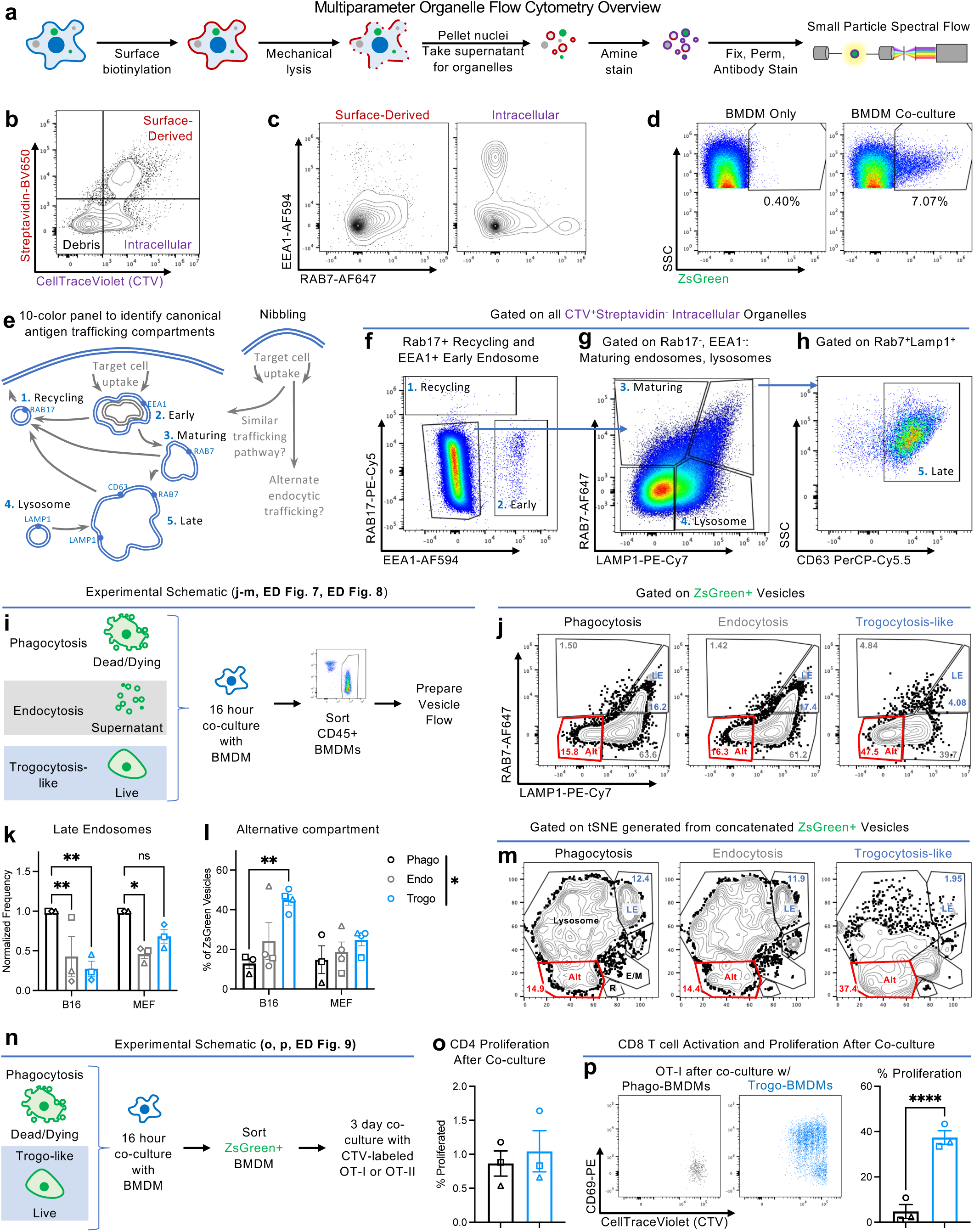
Multiparameter intracellular vesicle flow cytometry reveals that live-cell associated protein fills a discrete vesicular compartment. **a**, Cell surface membrane is biotinylated prior to mechanical lysis and low-speed centrifugation. Supernatant containing organelles are stained with CellTraceViolet, fixed, permed, and stained with antibodies against vesicle-associated antigens. **b,** Discrimination of Strepavidin^-^ CellTraceViolet^+^ intracellular, protein-containing particles. **c,** Vesicle-staining antibodies primarily stain intracellular particles. **d,** Discrimination of ZsGreen+ vesicles allows for tracking of vesicles containing target-cell-derived protein. **e-h,** Identification of vesicle compartments by vesicle flow panel. **i,** Experimental schematic to evaluate trafficking downstream of different sampling mechanisms. **j-m,** Representative flow plots (**j**) or tSNE (**m**) of ZsGreen^+^ vesicles derived from different sampling mechanism with summary quantification of proportion of ZsGreen^+^ vesicles in late endosome (**k**) and alternative compartment (**l**). **n,** Experimental schematic to evaluate T cell activation downstream of different sampling mechanisms. ZsGreen^+^ BMDMs were sorted after co-culture with live or apoptotic B16F10-ZsGreen-minOVA target cells and subsequently co-cultured with CellTraceViolet (CTV)-labeled OT-I CD8+ or OT-II CD4+ T cells in a 1:3 BMDM:T ratio for 3 days. **o,** Proliferation of OT-II CD4 T cells as read out by loss of CellTraceViolet after 3 days of co-culture with ZsGreen+ BMDMs that ingested material using phagocytosis (black) or trogocytosis-like (blue) mechanisms. **p,** Activation and proliferation of OT-I CD8 T cells as read out by CD69 expression or loss of CellTraceViolet after 3 days of co-culture with ZsGreen+ BMDMs from phagocytosis (black) or trogocytosis (blue) conditions. n = 3-4 biological replicates. Shown are mean of technical replicates +/- s.e.m. Two-way ANOVA with Dunnett’s multiple comparisons test or two-sided paired t-test. * p < 0.05, ** p < 0.01, **** p <0.0001. LE, late endosome; Alt, alternative compartment; E/M, early/maturing endosomes; R, recycling endosomes.

Vesicles derived from internalization are understood to flow through a pathway consisting of progressively more degradative compartments; early endosomes mature into late endosomes that fuse with lysosomes to form phagolysosomes^10,25^. We tuned our panel to identify known vesicular compartments, including RAB17^+^ recycling endosomes, EEA1^+^ early endosomes, LAMP1^+^ lysosomes, RAB7^+^ maturing endosomes, and RAB7^+^LAMP1^+^CD63^+^ late endosomes (**Fig. 4e-h**). By further incorporating antibodies recognizing antigen presentation molecules major histocompatibility complex (MHC) I and II, we could identify compartments consistent with antigen-loading. Consistent with previous reports^10^, MHC I is primarily excluded from LAMP1^+^ vesicles whereas MHC II can be detected across multiple vesicular compartments (**Extended Data 7a-b**). Together, these results highlighted the suitability of a high-dimensional vesicle analytic method to study composition and identity of vesicular compartments.

We then applied this organelle immunophenotyping method to compare the intracellular trafficking of ZsGreen^+^ antigen-containing vesicles after trogocytosis-like live-sampling to other sampling mechanisms, including endocytosis and phagocytosis (**Fig. 4i**). While endocytosis of soluble material derived from supernatant resulted in very low levels of ZsGreen uptake, sufficient ZsGreen vesicles were nevertheless detectable for downstream analysis (**Extended Data Fig. 7c-d**). In a conventional gating analysis, both phagocytosis and endocytosis resulted in distributions of proteins across the vesicle maturation spectrum, but revealed decreased trafficking of vesicles resulting from trogocytosis-like sampling to the late endosome (**Fig. 4j,k, Extended Data Fig. 7e-f**) and target cell-dependent changes in the co-localization of these with MHC-I but not MHC II (**Extended Data Fig. 7g**). Notably, while ∼90% of ZsGreen^+^ vesicles resulting from phagocytosis were associated with a conventional vesicle marker, a substantial portion of ZsGreen+ vesicles resulting from live-sampling did not associate with these canonical endocytic markers (**Fig. 4l**). This suggested the sequestration of live-sampled material into an alternative endocytic trafficking pathway, rather than those associated with degradation.

To analyze vesicle populations based on their intensity for each of our markers and in a way unbiased by previous classifications, we displayed the concatenated ZsGreen^+^ vesicles in a t-distributed stochastic neighbor embedding (tSNE) plot using only the vesicle protein marker intensities as parameters (**Extended Data Fig. 8a,b**). This demonstrated that while vesicles derived from each sampling method overlapped, there was a distinct shift in the distribution of vesicles comparing live-sampling (trogocytosis-like), to conventional phagocytosis of apoptotic cells (**Fig. 4m, Extended Data Fig. 8b**). When we overlayed intensities of markers such as LAMP1 and consistent with the conventional flow gating strategy, vesicles resulting from live-sampling were relatively depleted in late endosomes and enriched in an alternative vesicular compartment (**Fig. 4m, Extended Data Fig. 8c**).

To independently assess lysosomal delivery, we performed confocal microscopy and quantitative co-localization analysis. This confirmed that ZsGreen acquired through live-sampling showed significantly lower overlap with LAMP1 compared to antigen acquired from phagocytosis (**Extended Data Fig. 8d-f**), further supporting that live-cell-derived antigen has a unique vesicular fate.

## Live-sampling biases T cell activation by BMDMs

Immunologically, the nature of sampling may have repercussion for how internalized antigens are presented to T cells on Class I (‘cross-presentation’) versus Class II MHC. To examine how sampling through this trogocytosis-like mechanism affects antigen presentation, we isolated ZsGreen^+^ BMDMs after co-culture with DMSO- or Staurosporine-treated B16F10 cell expressing ZsGreen fused to the OT-I and OT-II ovalbumin (OVA) peptides (B16-ZsGminOVA)^13^ (**Fig. 4n, Extended Data Fig. 9a**). We detected only weak and indistinguishable activation in OT-II CD4 T cell activation after co-culture with BMDMs that had acquired antigen through phagocytosis (“Phago-BMDMs”) compared to those that had sampled live cells (“Trogo-BMDMs”) (**Fig. 4o, Extended Data Fig. 9b**). In contrast, Trogo-BMDMs but not Phago-BMDMs induced significant OT-I CD8 T cell proximal activation (CD69 upregulation) and proliferation (**Fig. 4p, Extended Data Fig. 9c**). This is consistent with the absence of trafficking of live-sampled material to the degradative late endosome and lysosomal compartments and suggests a bias for cross-presentation.

Trogocytosis resulting in trans-cellular membrane transfer (cross-dressing) and T cell activation has been described in dendritic cells^26^. To determine if this is occurring in our system, we performed antigen transfer and T cell stimulation assays using Balb/c-derived BMDMs (**Extended Data Fig. 9d**). MHC-I allele H2Kb was undetectable on either Balb/c BMDMs cultured alone or ZsGreen^+^ Balb/c BMDMs that had ingested material from B16-ZsGreen-minOVA target cells (**Extended Data Fig. 9e**). Consistent with this, ZsGreen^+^ Balb/c BMDMs induced significantly less antigen-specific CD8 T cell proliferation compared to ZsGreen^+^ B6 controls (**Extended Data Fig. 9f**). Thus, while limited peptide-MHC transfer may occur, the majority of CD8 T cell activation must be due to BMDM processing and presentation of ingested cytosolic antigen.

## Redirecting antigen to the lysosome is associated with decreased T cell activation

Material ingested through clathrin-, dynamin-independent mediated endocytosis is diverted away from lysosomal degradation^27^, and the SNX27-retromer complex can prevent lysosomal delivery of ingested material^28^ (**Fig. 5a**). Because live-sampling in our system is likewise dynamin-independent and leads to non-degradative trafficking, we examined how disrupting SNX27 affects antigen trafficking and ensuing T cell responses. CRISPR-Cas9-mediated targeting of the *Snx27* locus in BMDMs introduced indels and decreased protein expression at high efficiency (**Fig. 5b,c**). SNX27 KO did not impair ZsGreen sampling (**Extended Data Fig. 10a**). However, compared to *Rosa26*-targeted controls, SNX27 KO increased ZsGreen trafficking to LAMP1^+^ lysosomes and reduced the fraction of ZsGreen in the alternative compartment (**Fig. 5d,e, Extended Data Fig. 10b**), consistent with live-sampled material being diverted from the lysosome downstream of live-sampling.

**Figure 5:**
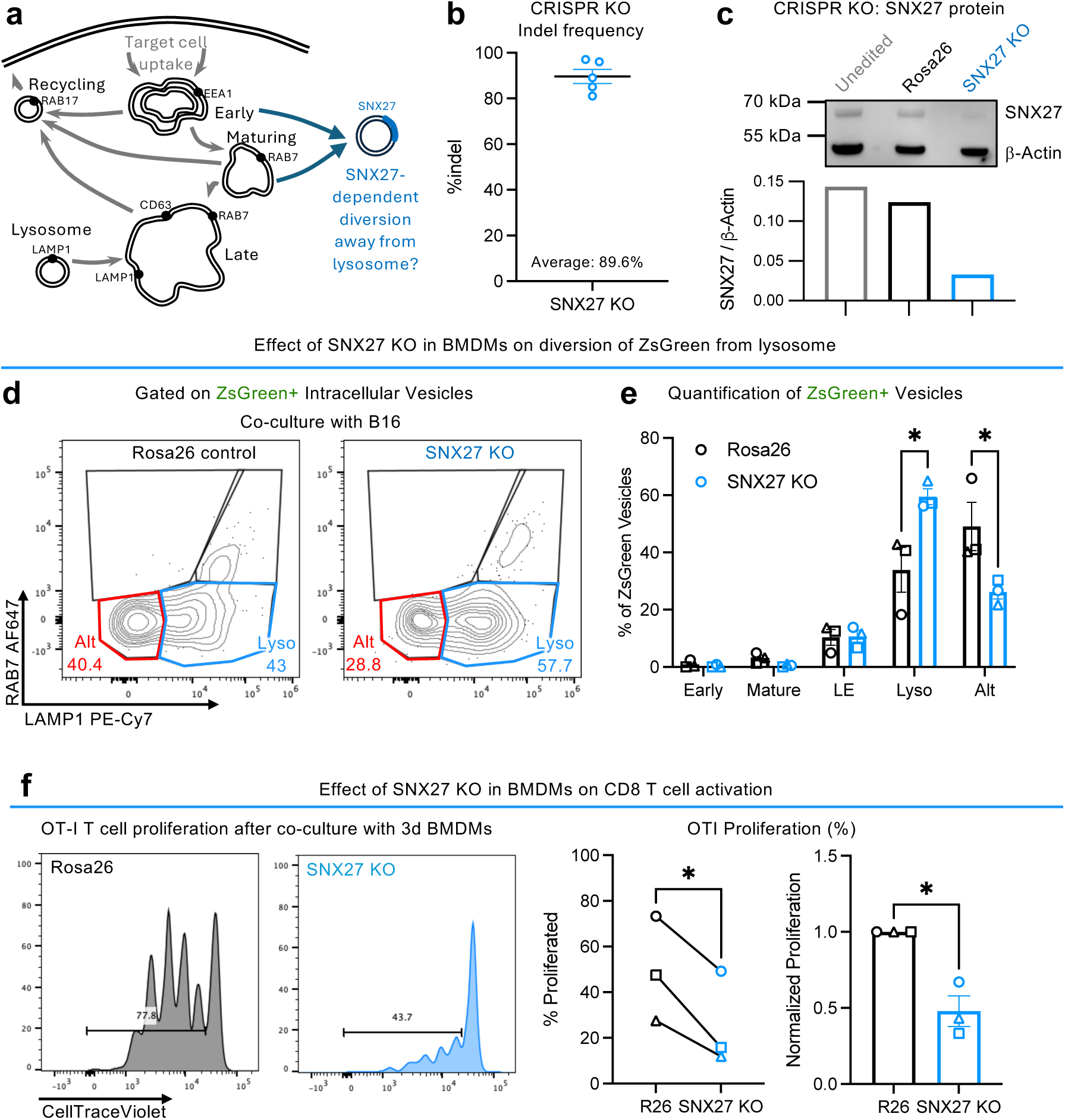
SNX27 KO in macrophages increases antigen trafficking to lysosomes and decreases CD8 T cell activation. **a**, Schematic illustrating SNX27-dependent diversion of material away from lysosomal maturation into alternative endocytic pathways. **b-c**, Validation of SNX27 KO in BMDMs by (**b**) indel quantification by ICE analysis (mean +/- s.d.) and (**c)** western blot (representative of 2 experiments). β-Actin was probed on the same gel as a loading control. For gel source data, see **Supplementary Fig. 1. d-e**, CRISPR-Cas9-targeted BMDMs were co-cultured with ZsGreen target cells to evaluate ZsGreen trafficking. **f**, OT-I CD8 T cell proliferation after 3-day co-culture with FACS-isolated ZsGreen^+^ *Rosa26*-targeted and SNX27 KO BMDMs that live-sampled B16-ZsGreen-minOVA target cells. n = 3-5 biological replicates. Shown are Shown are mean +/- s.e.m unless otherwise specified. Two-tailed paired t-test or two-way ANOVA with Sidak’s multiple comparison. *, p < 0.05

To determine whether these trafficking changes were associated with loss of cross-presentation, we co-cultured SNX27 KO or control BMDMs with B16-ZsGreen-minOVA target cells and isolated ZsGreen+ BMDMs for co-culture with OT-I CD8 T cells. Consistent with increased antigen degradation, SNX27 KO BMDMs induced less T cell proliferation (**Fig. 5f**). Importantly, SNX27 KO did not impair T cell proliferation when BMDMs were pulsed with SL8 peptide (**Extended Data Fig. 10c**), confirming that the effect is due to altered antigen processing rather than general defects in APC functionality.

## DISCUSSION

Together, these findings demonstrate a substantial sampling of live cells that takes place via a trogocytosis-like mechanism and which fills a distinct vesicular compartment that are particularly underrepresented in late endosomes^10^. That macrophages can constitutively ingest small amounts of material from live cells is in itself unsurprising; indeed a pruning function for macrophages has been demonstrated to be important to activate stem cells^29^ and to optimize brain circuitry via small synaptic pruning^30^. A key distinction here is in revealing that a very low-volume trogocytosis-like sampling may be ongoing at steady state in a variety of cells and may be a significant source of self-antigen from healthy cells, giving macrophages a distinct endocytic trafficking mechanism with which to present healthy-self antigens, notably to CD8 T cells. We also note a single previous suggestion of trogocytosis by dendritic cells which focused on a more conventional ‘ripping’ mechanism leading to direct transfer of pre-formed peptide-MHC from the donor cell membrane^26^. In contrast, using previously unavailable vesicle cytometry, our studies highlight the maintenance of sampled material in a depot that can be processed by the APC for presentation. A more stable depot of self-antigens provides a means of integrating and presenting the self-identity of many sampled cells, perhaps over days or longer. The longer retention of material sampled by this trogocytosis-like mechanism may contribute to a transferable pool of these vesicles, as have been observed to be exchanged amongst myeloid cells^13^. While we use model antigen and transgenic T cells in our study to reveal the capability of macrophages to cross-present live-cell-associated antigen by stimulating T proliferation, the functional role of this cross-presentation for maintaining or tuning homeostatic T cell populations that have undergone thymic self-tolerance requires further investigation. Ultimately, this work provides a novel framework to understand how the immune system can accumulate information about our healthy-self: a system that then can tune T cells to the normal concentrations of self-protein but may also be exploited by tumor cells to promote pathologic outcomes.

## Acknowledgments

We thank all members of the Krummel lab, Jayanta Debnath, Caleb R. Glassman, and Graham Barlow for discussion and support. We acknowledge the UCSF Parnassus Flow Core CoLabs, particularly Vinh Nguyen and Tamara Roach, for assistance generating flow cytometry data (RRID: SCR_018206; DRC Center Grant NIH P30 DK063720) and the Biological Imaging Development Colab with assistance in data collection, instrumentation, and processing (NIH shared equipment grant S10OD028611-01). pCDH-puro-CMV-GC3AI was a gift from Binghui Li (Addgene plasmid # 78910; http://n2t.net/addgene:78910; RRID:Addgene_78910). FcRγ KO mice were a gift from Dr. Oscar Aguilar. We thank Tristan Courau for providing the custom anti-TREM2 antibody. This work was supported by R01 AI175277-01 and 2 R01 CA197363-06A1 to MFK. A.C.F. was supported by T32HL007185 and the A.P. Giannini Postdoctoral Research Fellowship and Leadership Award. R.T. was supported by T32AI007334. N.S. was supposed by the Human Frontier Science Program (LT000061/2018-L). V.S.V. was supported by the UCSF Medical Scientist Training Program (T32GM141323) and the Achievement Rewards for College Scientists scholarship. M.K.R was supported by the Cancer Research Institute Irvington Postdoctoral Fellowship.

## Author contributions

A.C.F. and M.F.K. conceived the study, with N.S. contributing to initial conception. Methodology was developed by A.C.F., with N.S., and M.K.R. contributing to the initial methodological framework. Investigation was performed by A.C.F. (antigen transfer assays, imaging collection, imaging analysis, inhibitor studies, vesicle flow cytometry, T cell co-culture assays, indel analysis), R.T. (antigen transfer assays, imaging analysis, T cell co-culture assays), N.S. (in vivo experiments, imaging collection), V.S.V. (indel analysis, western blot, T cell co-culture assays), K.M. (imaging collection), M.K.R. (in vivo experiments, imaging collection), L.L. (antigen transfer assays), G.J. (cell isolation), and A.E (imaging analysis). Formal analysis was carried out by A.C.F. and R.T. Data curation was performed by A.C.F. Visualization was carried out by A.C.F., R.T., and N.S. A.C.F. wrote the original draft, and R.T. and M.F.K. reviewed and edited the manuscript. M.F.K. supervised the study and acquired funding.

## Declaration of Interests

Authors declare no competing interests.

Supplementary Information is available for this paper.

## METHODS

### RESOURCE AVAILABILITY

#### Lead Contact

Further information and requests should be directed to the Lead Contact: Matthew F. Krummel.

#### Materials Availability

Requests for resources and reagents should be directed to and will be fulfilled by the Lead Contact: Matthew F. Krummel.

### EXPERIMENTAL MODEL AND SUBJECT DETAILS

#### Mice

Mice were housed and bred under specific pathogen-free conditions at the University of California, San Francisco Laboratory Animal Research Center and all experiments conformed to ethical principles and guidelines approved by the UCSF Institutional Animal Care and Use Committee, the National Institutes of Health and the American Association of Laboratory Animal Care. C57BL/6 (RRID: MGI:2159769), Ai16^31^ (“mTmG”), B6;129P2-Fcer1g^tm1Rav^/J^32^ (“FcRγ KO”, RRID: MGI:2162818), and Balb/c (RRID: MGI:2161072) mice were purchased from Jackson Laboratory or bred in house. Both male and female mice ranging in age from 6 to 20 weeks were used for experimentation. Food and water were provided ad libitum.

To generate ZsGreen reporter mice, Ai6 mice were crossbred with K14-Cre, Scgb1a1-Cre-ERT2, Villin-Cre, B-actin-Cre mice. To induce ZsGreen expression in Scgb1a1-Cre-ERT2;Ai6 mice, mice were fed tamoxifen-containing chow ad libitum for two weeks.

For tumor studies, B16-F10 melanoma cancer cells were resuspended in PBS and mixed at a1:1 (v:v) ratio with growth factor reduced Matrigel Matrix (BD Biosciences) and 100000 cells in 50 uL were transplanted into the subcutaneous region of the mouse flank. On day 14 after tumor challenge, when tumors reached a size/volume of approximately 0.5 cm^3^, mice were sacrificed, tumors were excised and processed for downstream analysis.

#### Tumor cell lines

B16-F10 (ATCC, CRL-6475) was purchased. B16ZsGreen and B16ZsGreen minOVA cell lines were described previously^13^. In brief, to make this cell line, B16-F10 melanoma parental cells were genetically engineered using viral transduction with a ZsGreen or ZsGreen-minOVA construct. B16-GC3AI cell lines were genetically engineered to stably express the GFP-Casp3-activity-indicator using viral transduction with AddGene construct (Addgene #78910). Adherent cell lines were cultured at 37° C in 5% CO_2_ in DMEM (Invitrogen), 10% FCS (Benchmark), Pen/Strep/Glut containing 2 mM L-glutamine, 100 U/ml penicillin, and 100 mg/ml streptomycin (Invitrogen).

#### Mouse embryonic fibroblasts

Mouse embryonic fibroblasts (MEFs) were derived from Ai6;B-actinCre mice that were generated by crossing B-actin-Cre (strain # 033984, The Jackson Laboratory) to Ai6 mice (strain # 007906,The Jackson Laboratory) as previously described^33^. Briefly, day 13.5 embryos were collected from pregnant females, were minced, and digested with Trypsin. The retrieved cells were washed and plated in DMEM (Invitrogen), 15% FCS (Benchmark), Pen/Strep/Glut (Invitrogen) for overnight culture at 37° C in 5% CO_2_. Media was aspirated after 24 h to remove any cells remaining in suspension and replaced with fresh media. Cells were then grown to 70%–80% confluency and cryopreserved.

### METHOD DETAILS

#### Tissue digest and flow cytometry staining

##### Lung

Lungs were collected from mice following euthanasia by overdose with 2.5% Avertin. Lungs were placed in 3 mL of DMEM (Gibco) in C-Tubes (Miltenyi) briefly processed with a GentleMACS Dissociator (Miltenyi). 2 mL of DMEM with 0.26 U/mL LiberaseTM (Roche) and 0.25 mg/mLDNase I (Roche) was subsequently added and samples were then incubated at 37° C in a shaker for 30 min and dissociated to single cell suspensions by GentleMACS. Tissue homogenate was then passed through a 100 mm filter. Red blood cells were lysed with 3 mL of RBC Lysis Buffer (155 mM NH4Cl, 12 mM NaHCO3, 0.1 mM EDTA) per lung for 5 min at room temperature. Samples were then washed with FACS buffer (2% FBS, 2mM EDTA in PBS) and resuspended in appropriate buffer for staining for flow cytometry or FACS.

##### Tumor

For tumor digests, tumors from mice were harvested 14 days after injection. Tumors were minced and incubated in digestion buffer (100 U/mL collagenase type I (Roche), 500 U/mL collagenase type IV (Roche), and 200 mg/mL DNAse I (Roche) in RPMI1640 (Gibco)) for 30 minutes on a shaker at 37 ° C. Digestion mixtures were then pipetted repeatedly, followed by a second 15-minute incubation at 37° C. Cells were quenched with RPMI-1640 (GIBCO) plus 10% FCS, washed with FACS buffer, and filtered through a 100-mm cell strainer before staining for flow cytometry.

##### Lymph Nodes

Inguinal lymph nodes (LN) were dissected from mice, cleaned of fat, and digested as previously described^14^. In brief, LN were pierced and torn with sharp forceps in 24-well plates and incubated for 15 min at 37° C in 1 ml digestion buffer (100 U/ml collagenase type I (Roche), 500 U/ml collagenase type IV (Roche), and 20 μg/ml DNAse I (Roche) in RPMI-1640 (GIBCO)). Cells were pipetted up and down repeatedly, followed by a second 15-minute incubation at 37° C. After digestion, LN were washed with RPMI-1640 (GIBCO) plus 10% FCS, washed with FACS buffer, and filtered through 70 μm Nytex filters before staining for flow cytometry.

#### Imaging Sample Preparation, Image Acquisition, and Image Analysis

##### Two photon imaging of mouse lung, skin, tumor, and gut slices

Imaging of lung, skin, tumor, and gut slices were performed using a custom-built two-photon setup equipped with two infrared lasers (MaiTai: Spectra Physics; Chameleon: Coherent). The Chameleon laser was set to 950nm for excitation of ZsGreen. Emitted light was detected using a 25x 1.2NA water lens (Zeiss) coupled to a 6-color detector array (custom; using Hamamatsu H9433MOD detectors). Emission filters used were blue 475/23, green 510/42, yellow 542/27, red 607/70, far red 675/67. The microscope was controlled by the MicroManager software suite, and time-lapse z stack images were acquired every 90 s with five-fold averaging and z-step of 4mm. Data analysis was performed with Imaris software (Bitplane). For lung slices, mice were euthanized by anesthetic overdose with 1 mL 2.5% Avertin and then intubated by tracheotomy with the sheath from an 18-gauge i.v. catheter. Lungs were subsequently inflated with 1 mL of 2% low melting agarose (BMA) in sterile PBS at 37C. Agarose was then solidified by flooding the chest cavity with 4C PBS. Inflated lungs were excised, and the left lobe was cut into 300mm sections using a vibratome. For skin, tumor, and gut slices, mice were euthanized using CO2, obstructing fat was removed, and tissue sections were embedded in 4% low-melting agarose in PBS prior to sectioning. Sections were mounted on plastic coverslips and imaged by two-photon microscopy at 37C in carbogen (5% CO2:95% O2)-perfused RPMI-1640 media (GIBCO, without Phenol Red) in a heated chamber.

##### Spinning Disk Confocal

Glass bottom 96 well plates were coated in fibronectin and washed as described above. BMDMs, sorted ZsGreen^+^ BMDMs after antigenic transfer, or unsorted antigenic transfer assays were imaged. Live imaging by spinning disk confocal was performed at 37°C with a 488 and 561 nm lasers with 40% and 50% laser powers, respectively. For LAMP1 staining, cells were fixed with 4% PFA at room temperature for 15 minutes, permeabilized with 0.5% saponin at room temperature for 15 minutes, incubated with blocking buffer (1% BSA, 0.1% saponin, 5% normal rat serum) at room temperature for 60 minutes, and incubated at LAMP1-AF647 in blocking buffer at 4C overnight. Fixed cells were imaged with 488 and 640 at 25% laser powers.

##### Nikon NSPARC

Glass bottom 96 well flat bottom plates were coated with 50ug/mL fibronectin in H_2_O at 37°C for 1 h, or at 4°C overnight, before use. Fibronectin coated wells were washed 2x with PBS prior to use. 8×10^3^ TdTomato^+^ BMDMs were plated with 1.2×10^4^ ZsGreen+ B16F10s or ZsGreen+ MEFs and spun at 1500g for 5 min. Cells were incubated at 37°C for 2 hours prior to imaging. NSPARC imaging was performed at 37°C with 488 and 560 nm lasers with 1.0% and 2.0% laser powers, respectively.

##### Lattice light-sheet microscopy

5 mm round coverslips were cleaned by a plasma cleaner and coated with 2 μg/ml fibronectin in H_2_O at 37°C for 1 h, or at 4°C overnight, before use. Fibronectin coated coverslips were washed 2x with PBS prior to use. Cells were plated onto fibronectin-coated coverslips 20 min prior to imaging with a 10 min spin at 1,400 rpm and 4°C. Coverslip was immediately loaded into the sample bath with warmed imaging media and secured. Imaging was performed at 37°C with a 488 and 560 nm laser (MPBC). Exposure time was 10 ms per frame leading to a temporal resolution of ∼4.5. LLS microscope was a homebuilt clone of the scope described by Chen et al.^34^ with a Nikon CFI Apo LWD 25× W 1.1 NA 2 mm working distance objective, Hamamatsu Orca Flash 4.0 v2 camera, and Custom LabView acquisition software. Method was derived from previous work^19^.

##### Image Analysis

All computational image analysis was performed in Imaris version 9.9.1 or 10.2.0 (Bitplane) and Fiji version 2.16.0/1.54p. Particle area analysis was performed using the Analyze particles function in Fiji with the following parameters: size (.027- infinity) and circularity (0.00-1.00). Post processing raw data from lattice light-sheet images were deconvoluted using iterative Richardson-Lucy deconvolution as implemented in LLSpy. Briefly, images were deconvolved with a known point spread function that was recorded for each color prior to the experiment, as described previously^19^. A typical sample area underwent 15-20 iterations of deconvolution. For live imaging experiments, photobleaching correction was applied in FIJI using the histogram matching method. Particle volume was measured using the object-object statistics function in Imaris from the separated vesicle in frames 13-18.

For ZsGreen colocalization analysis with LAMP1, Mander’s coefficients were calculated using the JACoP plugin on ImageJ^35^. Otsu thresholding was used to identify optimal thresholding for the phagocytosis condition. To account for the low fluorescence intensities of ZsGreen vesicles in the live sampling condition, we opted for a manual threshold level of 550. Thresholding results were confirmed by comparing to ZsGreen negative BMDMs.

#### Flow cytometry and fluorescence-activated cell sorting

Zombie NIR Fixable Viability Kit (#423106; BioLegend), DAPI, or PI was used for exclusion of dead cells. Surface staining was performed with anti-mouse Fc receptor antibody (clone 2.4G2, UCSF Hybridoma Core) in FACS buffer for 30 min on ice. Supplementary Table 1 lists all antibodies referenced for flow cytometry and imaging experiments. Apoptotic cells were detected by staining with Annexin V AF647 (Biolegend #649012) and 1μg/mL DAPI in Annexin V binding buffer (Biolegend #422201). For all experiments involving intracellular staining, BD Cytofix/Cytoperm (#554722) was used. Following fixation and permeabilization, cells were incubated with Fc block for 10 min on ice prior to addition of intracellular stain. Flow cytometry was performed on a BD Fortessa instrument, and sorting was performed on BD FACSAria or BD FACSAria Fusion instruments. FlowJo software (BD Biosciences) was used for all analyses.

#### Generation of BMDMs

6-12 week old C57BL/6 mice were euthanized and their femurs and tibiae were excised. Bone marrow was crushed using a mortar and pestle. After pelleting the bone marrow, the red blood cells were lysed using RBC lysis buffer for 5 min at RT. Cells were washed with FACS buffer and filtered through a 100 mm cell strainer before being seeded at 1×106 cells / mL on a low-adherent cell culture dish in BMDM media (DMEM supplemented with 10% FBS (Benchmark), 50 mM beta-mercaptoethanol, Pen/Strup/Glut (Invitrogen), and 20 ng/ml M-CSF (Peprotech)). Fresh BMDM media was added on day 3-4 of culture. On day 6-7 of culture, BMDMs were harvested and used for experiments.

#### In vitro antigen transfer assay

BMDMs were isolated as described above and co-cultured with target B16-ZsGreen cells or MEF-ZsGreen cells at a 1:1 ratio for 16 hours before assays unless otherwise noted. Cells were plated in BMDM media in TC-treated 96 well flat-bottom plates prior to staining and analysis by flow cytometry.

#### Drug/Inhibitor antigenic transfer assay

Antigenic transfer assays were performed as described above. For all experiments wherein apoptosis was induced, B16F10 cells or MEFs were treated with 1μM Staurosporine (Tocris Bioscience, Cat# 1285) and washed with PBS prior to antigenic transfer assays. For exosome/microparticle inhibitory experiments, 5-(N,N-Dimethyl)amiloride hydrochloride (DMA, Sigma Aldrich, Cat# A4562) was plated at 0 h to a final concentration of 10μM. For endocytic inhibition experiments Dynole 34-2 (Cayman Chemical Company #34073) was plated at 0 h to a final concentration of 5μM. Caspase inhibition was performed using zVAD-FMK (Invivogen, Cat # tlrl-vad) at a final concentration of 20μM for B16-ZsGreen or 30μM for MEF-ZsGreen at 0 h. Signaling and cytoskeleton inhibition was performed using 10 μM PP1 (Cayman #14244), 5 μM Piceatannol (MedChemExpress #HY-13518), 10 μM GDC-0941 (Selleck Chemicals #S1065), 1.35 μM NAV-2729 (MedChemExpress #HY-112473), 25 μM NSC23766 (MedChemExpress #HY-15723), 10 μM ZCL278 (MedChemExpress #HY-13518), 200 μM CK-666 (Sigma-Aldrich #SML0006-5MG), or 5 μM SMIFH2 (MedChemExpress #HY-16931).

#### Drug/Inhibitor functional validation and cell toxicity validation

For exosome/microparticle inhibitor validation, target cells were cultured with DMA and supernatant was collected to quantify ZsGreen^+^ vesicles by small particle flow cytometry. For endocytosis inhibitor validation, exosomes were isolated from confluent B16-ZsGreen cultures using the ExoQuick kit (System Biosciences #EXOA5A-1), and BMDMs were cultured with exosomes and Dynole 34-2 or DMSO control. Exosome endocytosis was quantified with flow cytometry. For apoptosis inhibition studies, B16-ZsGreen or MEF-ZsGreen target cells were pre-treated with zVAD for 1 hour prior to Staurosporine treatment. %AnnexinV^+^DAPI^+/-^ apoptotic cells were evaluated by flow cytometry. For toxicity studies, cells were treated with drug or vehicle control for 16 hours and quantified by flow cytometry with CountBright beads (Invitrogen #C36950).

#### Supernatant/transwell antigenic transfer assay

Media was replaced on B16-ZsGreen cells or MEF-ZsGreen cells upon reaching 80% confluency. After 48 hours, supernatants were collected and used in antigenic transfer assays as described above. Transwell experiments were performed as described in antigen transfer assays with a 3μm pore transwell insert separating BMDM and ZsGreen+ target cells or supernatants.

#### Antibody Opsonization and Blockade

For opsonization experiments, B16-ZsGreen or MEF-ZsGreen target cells were pre-coated with 10ug/mL Armenian Hamster IgG (BD Pharmigen, Cat# 553969), 10μg/mL Rat IgG2ak (clone 2A3, invivoMab Cat# BE0089), 10μg/mL CD29 (eBioHmb1-1, eBioscience Cat #16-0291-85), 10μg/mL aCD98 (RL388, Biolegend Cat# 128202), normal mouse serum (Jackson ImmunoResearch Cat# 015-000-120), or 500μg/mL IgG from normal mouse serum at 37C for 30-60 minutes and washed with PBS prior to co-culture with BMDMs. IgG from normal mouse serum was isolated using Protein G dynabeads (Invitrogen Cat# 10003D) according to manufacturer instructions and dialyzed with PBS (Slide-A-Lyzer). Concentration of isolated IgG was quantified using IgG ELISA (abcam, Cat # ab151276).

For antibody blockade, BMDMs were pre-treated with 10μg/mL Rat IgG2b (clone MPC-11, invivoMab Cat# BE0086), CD16/32-blocking antibody (clone 2.4G2), CD11b-blocking antibody (clone M17/0, Biolegend Cat# 101202, RRID: AB_312785), CD49e blocking antibody (clone 5H10-27(MRF5), Biolegend Cat# 103801, RRID: AB_313050), SIRPα blocking antibody (Clone P84, Biolegend Cat# 144035, RRID: AB_2832516), or custom afucosylated mouse IgG2a monoclonal blocking antibody against TREM2 (provided by Tristan Courau, synthesized by Evitra) for 1 hour before 16-hour co-culture antigen transfer assays.

#### CRISPR Editing of Primary BMDMs

CRISPR editing of primary BMDMs were performed as previously described^36^. Briefly, sgRNA targeting *Rosa26 (ACUCCAGUCUUUCUAGAAGA)*, *Fcgr1* (*GAUCACCUUGCAGCCUCCAU*), *Fcgr3* (*UGGUGAAACUGGACCCCCCA*), *Fcgr4* (*GGUGAACCUAGACCCCAAGU*), *Itgam (GAAGCCAUGACACAAGGCUA)*, *Itgav*(*UUGAAUCAAACUCAAUGGGC*, *CCUGUUGAAUCAAACUCAAU*), *Tlr2* (*UUGGCUCUUCUGGAUCUUGG*), *C5ar1* (*CAUGGAUCCUAACAUACCUG*, *GAUCCUAACAUACCUGCGGA*, *AUGGCAUUCACCUCCCGAAG*), *Cd93*(*CAGGAACAAACCAGUUGAGA*, *AGAAGAAUGGCCAUCUCAAC*, *CUGGUUUGUUCCUGCUGCUG*), and *SNX27*(*GGAACGGCGUGAAUGUUGAG*, *UGAGGGGGCGACACACAAGC*, *GUGGUGGACCUGAUCCGAGC*) were ordered from Synthego or IDT and reconstituted to 100uM in TE (IDT). sgRNA and Cas9 (6.5mg/mL) was complexed in a 2:1 molar ratio at room temperature for at least 10 minutes. At 3 days after initial plating, BMDM differentiation cultures were harvested, and cells were washed with PBS prior to electroporation with ribonucleoprotein (RNP) containing gene-targeting sgRNA complexed in P3 primary cell solution with electroporation code CM137 using the Lonza 4D Nucleofector. After an additional 4 days of differentiation, gene-edited BMDMs were either stained with antibodies for validation of protein knockdown or co-cultured with ZsGreen target cells for 16 hours before evaluated by flow cytometry. For indel quantification with ICE, DNA was extracted with QuickExtract for PCR of the targeted locus with Phusion Plus Green PCR Master Mix (Thermo Scientific #F632S) and primers (5’ *CCCCATCTTTCCCACATGCT* 3’, 5’ *ATTACTGTAGGCCACCCCCT* 3’).

#### NicheNet analysis to predict receptors

To identify potential BMDM surface proteins mediating live-sampling using NicheNet^22^, we used the BMDM datasets from Johannessen, *et al*^37^, B16 dataset from Griffin, Wu, Iracheta-Vellve, *et al*^38^, and MEF dataset from Jaber, *e al*^39^. The BMDM was set as the “receiver” population and B16/MEF were set as sender populations. A threshold for expressed genes for each population was defined based on Gaussian fit of gene expression BMDM receptors were prioritized if predicted to interact with both cell types, overlap with known phagocytic modulators, and confirmed to be expressed at the protein level.

#### Vesicle Flow Cytometry

##### Sample Preparation and Sorting

16 hours prior to sorting, BMDMs were cultured with live target cells (“Trogocytosis”) in a 1:1 ratio, Staurosporine-treated target cells (“Phagocytosis”) in a 1:1 ratio, or supernatant derived from target cells (“Endocytosis”). Cells were harvested, washed with FACS buffer, stained with CD45-BUV395 with FcBlock for 30 minutes on ice, washed with FACS buffer, and resuspended in FACS buffer + 1 mg/mL PI at 10E6 cells/mL. CD45^+^ cells were sorted on FACSAria and FACSFusion machines on “Purity” into FACS buffer kept at 4C.

##### Surface Biotinylation and Lysis

Cells were washed 3x in PBS, resuspended in PBS at 25E6 cells/mL then biotinylated with 80 uL of 10mM EZ-Link Sulfo-NHS-SS-Biotin (ThermoFischer Scientific #21331) per mL of reaction volume for 30 minutes at 4C. Cells were washed 3x in PBS, resuspend in at least 500 μL homogenization buffer (250mM sucrose, 200mM PMSF, 250mM sucrose, 3mM Imidazole, 1X protease inhibitor filtered through 0.1um filter) at a concentration of 10E6 cells/mL, and drawn 15 times up and down through a 22G needle. Cell homogenate was spun for 4 minutes at 150xg, and post-nuclear supernatant was spun at 3000xg for 5 min at 4C to collect vesicles.

##### Staining

All staining reagents were filtered using a 0.1μm filter prior to use and vesicles were pelleted using spin speeds of 3000xg for 5 min at 4C for all steps. Vesicles were stained with 50μM CellTrace Violet (ThermoFischer Scientific Cat# C34557) for 5 minutes at room temperature and washed once with PBS. Samples were fixed for 20 minutes at room temperature with fixation buffer (Invitrogen Cat# 88-8824-00), washed with permeabilization buffer (Invitrogen Cat# 88-8824-00) twice, and stained in 50 μL of permeabilization buffer with 1% FCS with FcBlock for 30 minutes at room temperature. Vesicles were washed with permeabilization buffer and resuspend in 100 μL PBS for acquisition.

##### Sample Acquisition and Analysis

Aurora Cytek flow cells were soaked with contrad overnight with long clean prior to acquisition with filtered reagents. Calibration beads and samples were acquired on the Aurora Cytek with Enhanced Small Particle detection.

#### T cell stimulation assays

ZsGreen^+^ BMDMs were sorted from BMDM 16-hour co-cultures with DMSO- or Staurosporine-pretreated B16zsGreenminOVA. To induce MHC II expression on BMDMs, sorted cells were stimulated with 20ng/mL IFNγ for 2 hours and then washed prior to co-culture with T cells. OTI and OTII T cells were isolated from spleens of TCR transgenic mice after RBC lysis using either a CD8 or CD4 EasySep enrichment kit (STEMCELL Technologies), respectively. T cells were labeled with CellTraceViolet (ThermoFischer Scientific) at 37C in PBS for 15 minutes and washed in RPMI prior to use. T cell stimulation assay was performed as previous described. Briefly, sorted BMDMs and T cells were added to the wells of a 96-well V-bottom plate at a 1:3 ratio in RPMI (Gibco) supplemented with 10% FCS (Benchmark), Pen/Strep/Glut (Invitrogen), and 50 mM beta-mercaptoethanol (Thermo Fisher). Cells were harvested for analysis 3 days later. Dilution of cell permeable dye CellTraceViolet and expression of CD69 were used as indicators of T cell stimulation.

#### Western Blot

Whole-cell protein lysates were obtained from BMDMs in Pierce RIPA buffer [Thermo Scientific, Cat# 89901] and ProteoGuard EDTA-Free Protease Inhibitor Cocktail (Takara, Cat# 635673). Lysates were extracted via sample agitation at 4°C for 30 minutes, followed by 4°C centrifugation at 16,000 RPM for 20 minutes. Protein concentration was determined with Pierce BCA Protein Assay Kit (Thermo Fisher). Lysates were denatured in NuPage LDS sample buffer (Invitrogen, Cat# NP0007) supplemented with NuPAGE Sample Reducing Agent (Invitrogen, Cat# NP0009) by incubation at 95°C for 5 minutes. Denatured samples were loaded onto a NuPAGE 10% (Cat# NP0302BOX) or 4% to 12% Bis-Tris (Cat# NP0323BOX) polyacrylamide gel (Thermo Fisher) with PageRuler Plus Prestained Protein Ladder (Thermo Scientific, Cat#26619) and electrophoresed in MES running buffer (Invitrogen). Gels were transferred to PVDF membranes with the iBlot 2 system (Invitrogen, Cat# IB21001), blocked with Pierce Clear Milk Blocking Buffer (Thermo Scientific, Cat# 37587), and incubated with primary antibodies according to the manufacturers’ directions. Primary antibodies, SNX27 (Abcam, Clone: EPR218130-16, Cat# ab315897), and β-Actin (Invitrogen, Cat #PA1-183, RRID: AB_2539914) were detected with Goat Anti-Rabbit IgG HRP-conjugated secondary antibodies (Southern Biotech, Cat# 4050-05, RRID: AB_2795955) and SuperSignal West Femot Maximum Sensitivity Substrate (Thermo Scientific, #34095). Membranes were imaged with the Licor Odyssey XF system.

### QUANTIFICATION AND STATISTICAL ANALYSIS

#### Statistical analysis

Unless specifically noted, all data are representative of >3 separate experiments. Experimental group assignment was determined by genotype or, if all WT mice, by random designation. Statistical analyses were performed using GraphPad Prism software. Error bars represent standard error of the mean (S.E.M.) calculated using Prism, and are derived from triplicate or greater experimental conditions. Specific statistical tests used were paired and unpaired t-tests and p values <0.05 were considered statistically significant. For pairwise comparisons, unpaired t tests were used unless otherwise noted. For statistical measures between more than two groups, one-way ANOVA would be performed unless otherwise noted. Investigators were not blinded to group assignment during experimental procedures or analysis.

### DATA AVAILABILITY

The data reported are in the Article and its associated Supplementary, and all relevant data are available from the corresponding author on request. Source data are provided with this paper. Transcriptomic data were downloaded from NCBI GEO database, accession numbers GSE99759, GSE155972, and GSE171127.

## Extended Data Figure Legends

**Extended Data Figure 1: ZsGreen uptake from tissues is not due to Cre mis-expression.**

**a**, Gut taken from mice expressing ZsGreen under the Villin promoter. Isolation of myeloid cells from Villin-Cre;ZsGreen mice shows uptake in gut in macrophages as well as DCs. **b**, Representative flow (left) or quantification of ZsGreen (right) in lung myeloid cell populations in B6 wildtype mice, Scgb1a1-CreERT2;ZsGreen mice with lung-specific ZsGreen expression, and B-actin-Cre-ZsGreen mice. n = 2-6 mice per condition. **c**, Bone marrow transplant of B6 wildtype mice into B-actin-Cre;ZsGreen expressing mice. Myeloid cells isolated from skin-draining inguinal lymph nodes show ZsGreen sampling. **d**, Intracellular staining for airway specific PLVAP protein in lung (black) compared to splenic (gray) myeloid cell populations with ZsGreen+ lung macrophages (green) enriching for PLVAP signal. Shown are mean +/- standard deviation. Representative of 3 experiments, n = 3-6 mice per experiment. M, macrophage; Neu, neutrophil; Mo, monocyte; mDC2, migratory cDC2; mDC1, migratory cDC1; rDC1, residential cDC1; rDC2, residential cDC2; MoMac, Monocyte/Macrophage.

**Extended Data Figure 2: Live cells can be sampled in a cell-contact dependent manner without caspase activation**

**a**, Visualization of ZsGreen (C1) puncta ingested from target cells within membrane Tomato (C2) labeled BMDMs. ZsGreen+ BMDMs were sorted and imaged after co-culture with B16-ZsGreen cells. **b**, BMDMs were co-cultured directly with MEFs, with MEF supernatant, or with a transwell insert containing MEFs. **c-d**, Target cells (**c**) or BMDM-MEF co-cultures (**d**) were treated with DMA or DMSO vehicle control. Extracellular vesicle production in supernatant (**c**) or ZsGreen uptake (**d**) was measured by flow cytometry. **e**, Effect of Dynole on endocytosis of ZsGreen^+^ exosomes by BMDMs. **f**, BMDM-MEF co-cultures were treated with indicated inhibitors or DMSO vehicle control. **g**, Target cells were treated with staurosporine and/or zVAD and AnnexinV^+^DAPI^+/-^ apoptotic cells were measured with flow cytometry. **h**, BMDM co-culture with B16-F10 expressing GFP caspase 3 activity indicator (B16-GC3AI) demonstrates uptake is predominantly from live cells. n = 3 biological replicates. Shown are mean of technical replicates +/- s.e.m. Paired t-test or one-way ANOVA. * p < 0.05, ** p < 0.01.

**Extended Data Figure 3: A trogocytosis-like process can sample target cell protein**

**a**, Three-dimensional surface renderings from the YZ (left) and XY (bottom right) directions to identify intracellular localization of separated vesicle from same ROI as Fig. 3a, b. **b**, Two-color volume rendering showing interactions between TdTomato+ BMDM and ZsGreen+ B16(left) captured immediately after the series in Fig 3b. Three-dimensional surface renderings(right) of ROIs show separated vesicle trafficking towards the nucleus. Time shown as hh:mm:ss. **c**, Spinning-disk confocal imaging of ZsGreen+ B16 and TdTomato+ BMDM showing ingested ZsGreen+ puncta within BMDMs. Time shown as hh:mm. **d**, Three-dimensional surface rendering from Lattice light-sheet imaging of TdTomato+ BMDM and ZsGreen+ B16

**Extended Data Figure 4: Antibody opsonization amplifies live-cell sampling**

**a**, Schematic outlining methods used to block or knock-out (KO) FcγR binding and signaling. **b**, BMDMs targeted at the Rosa26 control locus or receptor loci using CRISPR-Cas9 RNP. **c**, BMDMs were pre-incubated with 10 ug/mL 2.4G2 CD16/32 blocking antibody or Rat IgG2b prior to co-culture with antibody-opsonized B16-ZsGreen target cells. **d-e**, CRISPR-targeted BMDMs (d) or BMDMs isolated from B6 wildtype or FcγR KO (e) mice were co-cultured with B16-ZsGreen or MEF-ZsGreen target cells pre-coated with antibody. **f-g**, BMDMs co-cultured with B16-ZsGreen pre-coated with serum (f), or IgG isolated from mouse serum (g). n = 3 biological replicates. Shown are mean of technical replicates +/- s.e.m. Paired t-test. * p < 0.05, ** p < 0.01.

**Extended Data Figure 5: Knockout of CD11b and CD93 decreases live sampling**

**a**, Representative protein knockdown 4 days after CRISPR/Cas9-mediated gene targeting. **b**, ZsGreen uptake in KO cells compared to Rosa26 (R26) controls. **c**, BMDMs were pre-treated with 1ug/uL IgG2b or CD11b-blocking antibody for 1 hour before 16-hour co-culture antigen transfer assays. **d**, Schematic outlining strategy to identify potential modulators of live sampling. **e**, BMDMs were co-cultured with B16-ZsGreen or MEF-ZsGreen target cells for 16 hours with 5 ug/mL blocking antibody or the appropriate isotype control prior to flow cytometric analysis of ZsGreen uptake. f-g BMDMs targeted at the *Rosa26* control locus or receptor loci using CRISPR-Cas9 RNP and co-cultured with B16-ZsGreen or MEF-ZsGreen target cells for 16 hours prior to flow cytometric analysis of ZsGreen uptake (f) and receptor expression (g). Shown are mean +/- s.e.m. n = 2-3 experiments per receptor, 2-3 technical replicates per experiment. Kruskal-Wallis test with Dunn’s multiple comparisons test. * p < 0.05.

**Extended Data Fig. 6: Src, PI3K, and Arp2/3 inhibition decreases live sampling**

**a**, Schematic illustrating inhibitors used to target components of receptors-mediated uptake. **b-e**, BMDMs, B16-ZsGreen target cells, and MEF-ZsGreen target cells were cultured alone or co-culture for antigen transfer assay and treated with DMSO control, 10 uM PP1, 5 uM Piceatannol, 10 uM GDC-0941, 1.35 uM NAV-2729, 25 uM NSC23766, 10 uM ZCL278, 200 uM CK-666, or 5 uM SMIFH2 for 16 hours prior to flow cytometry analysis for cell number (**b**) or ZsGreen uptake (**c-e**). Data are normalized to DMSO control and compiled over 3-6 experiments. Shown are mean +/- s.e.m. Kruskal Wallis test and Dunn’s multiple comparisons. *, p < 0.05; **, p < 0.01.

**Extended Data Figure 7: Live-cell associated antigen fills a discrete vesicular compartment**

**a-b**, Distribution of MHC I (a) and MHC II (b) in identified vesicular compartments inclusive of both ZsGreen+ and ZsGreen- vesicles. n = 3-4 biological replicates. Shown are mean of technical replicates +/- s.e.m. **c**, Representative sorting gate and purity check for CD45+ BMDMs from 16-hour co-culture. **d**, Representative ZsGreen+ gating from different vesicle flow preps gated on CellTraceViolet+Streptavidin- intracellular vesicles. **e**, Representative flow cytometry of vesicle populations from phagocytosis, endocytosis, and trogocytosis of B16-ZsGreen or MEF-ZsGreen target cells. **f**, Summary quantification of proportion of ZsGreen+ vesicles in different vesicular compartments. **g**, Proportion of ZsGreen+ vesicles that were either MHC I+ or MHC II+. n = 3-4 biological replicates. Shown are mean of technical replicates +/- s.e.m.

**Extended Data Figure 8: Live-cell associated antigen fills a LAMP1-negative vesicular compartment a**, tSNE of concatenated ZsGreen+ vesicles from one experiment demonstrating gating of different vesicular compartments. **b**, Distribution of ZsGreen+ vesicles in derived from different sampling mechanism. **c**, Summary quantification of proportion of ZsGreen+ vesicles in late endosome and alternative vesicular compartment gates by gating on tSNE populations. n = 3-4 biological replicates. Shown are mean of technical replicates +/- s.e.m. **d-f**, Sorted ZsGreen+ BMDMs after co-culture with killed B16-ZsGreen (Phago-BMDM) and live B16-ZsGreen (Trogo-BMDM) were fix, permeabilized, and stained with LAMP1-AF647. Representative images are shown in (d,e) and quantification per field of view is shown in (f). Phago, n = 24; Trogo, n=96. Two-sided student’s t-test or Two-way ANOVA. * p < 0.05, ** p < 0.01.

**Extended Data Figure 9: Macrophages process live-cell associated antigen to affect CD8 T cell activation**

**a**, Representative flow plots showing sorting and purity of ZsGreen+ BMDMs after co-culture with live or apoptotic B16F10-ZsGreen-minOVA target cells. **b-c**, Activation and proliferation of OT-I CD8 T cells (**b**) and OT-II CD4 T cells (**c**) as read out by CD69 expression after 3 days of co-culture with ZsGreen+ BMDMs from phagocytosis (black) or trogocytosis (blue) conditions. **d**, Experimental schematic to evaluate strain specific MHC-I transfer and T cell activation by ZsGreen^+^ B6 or Balbc-derived BMDMs. **e**, H2-Kb expression on B16 ZsGreen-minOVA target cells; ZsGreen+ BMDMs after co-culture with B16 ZsGreen-minOVA cels, and BMDMs cultured alone. **f**, OT-I CD8 T cell proliferation after 3-day co-culture with FACS-isolated ZsGreen+ B6 or Balb/c BMDMs. n = 3 biological replicates with n = 3 technical replicates. Shown are mean of technical replicates +/- s.e.m. Two-sided paired t-test. * p < 0.05.

**Extended Data Fig. 10: SNX27 KO reduces antigen diversion from the lysosome but not uptake or presentation capacity**

**a-c**, ZsGreen uptake (a) and vesicle distribution (b,c) in SNX27 KO cells compared to Rosa26 (R26) controls. **d**, OT-I proliferation after 3-day co-culture with peptide-pulsed SNX27 KO and Rosa26 control BMDMs.

## Notes

### Competing Interest Statement

The authors have declared no competing interest.

### Summary of Updates

We have: 1. Added additional controls for inhibitors in Figure 2; 2. Used blocking antibodies and genetic KO to examine the contribution of surface proteins and signaling components to uptake; and 3. Revealed that SNX27 KO can re-direct live-sampled material to the lysosome and decrease T cell proliferation.

## REFERENCES

1. Medawar, P.B. (1961). IMMUNOLOGICAL TOLERANCE.

2. Goodnow, C.C., Sprent, J., de St Groth, B.F., and Vinuesa, C.G. (2005). Cellular and genetic mechanisms of self tolerance and autoimmunity. Nature 435, 590–597. 10.1038/nature03724.

3. Kamradt, T., and Mitchison, N.A. (2001). Tolerance and Autoimmunity. N. Engl. J. Med. 344, 655–664. 10.1056/NEJM200103013440907.

4. Kenison, J.E., Stevens, N.A., and Quintana, F.J. (2023). Therapeutic induction of antigen-specific immune tolerance. Nat. Rev. Immunol., 1–20. 10.1038/s41577-023-00970-x.

5. Cummings, R.J., Barbet, G., Bongers, G., Hartmann, B.M., Gettler, K., Muniz, L., Furtado, G.C., Cho, J., Lira, S.A., and Blander, J.M. (2016). Different tissue phagocytes sample apoptotic cells to direct distinct homeostasis programs. Nature 539, 565–569. 10.1038/nature20138.

6. Kushwah, R., Wu, J., Oliver, J.R., Jiang, G., Zhang, J., Siminovitch, K.A., and Hu, J. (2010). Uptake of apoptotic DC converts immature DC into tolerogenic DC that induce differentiation of Foxp3+ Treg. Eur. J. Immunol. 40, 1022–1035. 10.1002/eji.200939782.

7. Myers, D.R., Zikherman, J., and Roose, J.P. (2017). Tonic Signals: Why do Lymphocytes Bother? Trends Immunol. 38, 844–857. 10.1016/j.it.2017.06.010.

8. Stefanová, I., Dorfman, J.R., and Germain, R.N. (2002). Self-recognition promotes the foreign antigen sensitivity of naive T lymphocytes. Nature 420, 429–434. 10.1038/nature01146.

9. Hogquist, K.A., and Jameson, S.C. (2014). The self-obsession of T cells: how TCR signaling thresholds affect fate “decisions” and effector function. Nat. Immunol. 15, 815–823. 10.1038/ni.2938.

10. Blum, J.S., Wearsch, P.A., and Cresswell, P. (2013). Pathways of Antigen Processing. Annu. Rev. Immunol. 31, 443–473. 10.1146/annurev-immunol-032712-095910.

11. Greene, J.T., Brian, B.F., Senevirathne, S.E., and Freedman, T.S. (2021). Regulation of myeloid-cell activation. Curr. Opin. Immunol. 73, 34–42. 10.1016/j.coi.2021.09.004.

12. White, S.R. (2011). Apoptosis and the Airway Epithelium. J. Allergy 2011, 948406. 10.1155/2011/948406.

13. Ruhland, M.K., Roberts, E.W., Cai, E., Mujal, A.M., Marchuk, K., Beppler, C., Nam, D., Serwas, N.K., Binnewies, M., and Krummel, M.F. (2020). Visualizing Synaptic Transfer of Tumor Antigens among Dendritic Cells. Cancer Cell 37, 786–799.e5. 10.1016/j.ccell.2020.05.002.

14. Roberts, E.W., Broz, M.L., Binnewies, M., Headley, M.B., Nelson, A.E., Wolf, D.M., Kaisho, T., Bogunovic, D., Bhardwaj, N., and Krummel, M.F. (2016). Critical Role for CD103+/CD141+ Dendritic Cells Bearing CCR7 for Tumor Antigen Trafficking and Priming of T Cell Immunity in Melanoma. Cancer Cell 30, 324–336. 10.1016/j.ccell.2016.06.003.

15. Pham, T., Mero, P., and Booth, J.W. (2011). Dynamics of Macrophage Trogocytosis of Rituximab-Coated B Cells. PLOS ONE 6, e14498. 10.1371/journal.pone.0014498.

16. Blander, J.M. (2017). The many ways tissue phagocytes respond to dying cells. Immunol. Rev. 277, 158–173. 10.1111/imr.12537.

17. Yin, C., and Heit, B. (2021). Cellular Responses to the Efferocytosis of Apoptotic Cells. Front. Immunol. 12. 10.3389/fimmu.2021.631714.

18. Zhang, J., Wang, X., Cui, W., Wang, W., Zhang, H., Liu, L., Zhang, Z., Li, Z., Ying, G., Zhang, N., et al. (2013). Visualization of caspase-3-like activity in cells using a genetically encoded fluorescent biosensor activated by protein cleavage. Nat. Commun. 4, 2157. 10.1038/ncomms3157.

19. Cai, E., Marchuk, K., Beemiller, P., Beppler, C., Rubashkin, M.G., Weaver, V.M., Gérard, A., Liu, T.-L., Chen, B.-C., Betzig, E., et al. (2017). Visualizing dynamic microvillar search and stabilization during ligand detection by T cells. Science 356, eaal3118. 10.1126/science.aal3118.

20. Wang, Y., Krémer, V., Iannascoli, B., Goff, O.R.-L., Mancardi, D.A., Ramke, L., de Chaisemartin, L., Bruhns, P., and Jönsson, F. (2022). Specificity of mouse and human Fcgamma receptors and their polymorphic variants for IgG subclasses of different species. Eur. J. Immunol. 52, 753–759. 10.1002/eji.202149766.

21. Hong, S., Beja-Glasser, V.F., Nfonoyim, B.M., Frouin, A., Li, S., Ramakrishnan, S., Merry, K.M., Shi, Q., Rosenthal, A., Barres, B.A., et al. (2016). Complement and microglia mediate early synapse loss in Alzheimer mouse models. Science 352, 712–716. 10.1126/science.aad8373.

22. Browaeys, R., Saelens, W., and Saeys, Y. (2019). NicheNet: modeling intercellular communication by linking ligands to target genes. Nat. Methods. 10.1038/s41592-019-0667-5.

23. Cockram, T.O.J., Dundee, J.M., Popescu, A.S., and Brown, G.C. (2021). The Phagocytic Code Regulating Phagocytosis of Mammalian Cells. Front. Immunol. 12. 10.3389/fimmu.2021.629979.

24. Hoffmann, E., Pauwels, A.-M., Alloatti, A., Kotsias, F., and Amigorena, S. (2016). Analysis of Phagosomal Antigen Degradation by Flow Organellocytometry. BIO-Protoc. 6. 10.21769/BioProtoc.2014.

25. Song, W., Cho, H., Cheng, P., and Pierce, S.K. (1995). Entry of B cell antigen receptor and antigen into class II peptide-loading compartment is independent of receptor cross-linking. J. Immunol. 155, 4255–4263. 10.4049/jimmunol.155.9.4255.

26. Harshyne, L.A., Zimmer, M.I., Watkins, S.C., and Barratt-Boyes, S.M. (2003). A Role for Class A Scavenger Receptor in Dendritic Cell Nibbling from Live Cells. J. Immunol. 170, 2302–2309. 10.4049/jimmunol.170.5.2302.

27. Shafaq-Zadah, M., Dransart, E., and Johannes, L. (2020). Clathrin-independent endocytosis, retrograde trafficking, and cell polarity. Curr. Opin. Cell Biol. 65, 112–121. 10.1016/j.ceb.2020.05.009.

28. Burd, C., and Cullen, P.J. (2014). Retromer: A Master Conductor of Endosome Sorting. Cold Spring Harb. Perspect. Biol. 6, a016774. 10.1101/cshperspect.a016774.

29. Wattrus, S.J., Smith, M.L., Rodrigues, C.P., Hagedorn, E.J., Kim, J.W., Budnik, B., and Zon, L.I. (2022). Quality assurance of hematopoietic stem cells by macrophages determines stem cell clonality. Science 377, 1413–1419. 10.1126/science.abo4837.

30. Weinhard, L., di Bartolomei, G., Bolasco, G., Machado, P., Schieber, N.L., Neniskyte, U., Exiga, M., Vadisiute, A., Raggioli, A., Schertel, A., et al. (2018). Microglia remodel synapses by presynaptic trogocytosis and spine head filopodia induction. Nat. Commun. 9, 1228. 10.1038/s41467-018-03566-5.

31. Muzumdar, M.D., Tasic, B., Miyamichi, K., Li, L., and Luo, L. (2007). A global double-fluorescent Cre reporter mouse. genesis 45, 593–605. 10.1002/dvg.20335.

32. Takai, T., Li, M., Sylvestre, D., Clynes, R., and Ravetch, J.V. (1994). FcR γ chain deletion results in pleiotrophic effector cell defects. Cell 76, 519–529. 10.1016/0092-8674(94)90115-5.

33. Headley, M.B., Bins, A., Nip, A., Roberts, E.W., Looney, M.R., Gerard, A., and Krummel, M.F. (2016). Visualization of immediate immune responses to pioneer metastatic cells in the lung. Nature 531, 513–517. 10.1038/nature16985.

34. Chen, B.-C., Legant, W.R., Wang, K., Shao, L., Milkie, D.E., Davidson, M.W., Janetopoulos, C., Wu, X.S., Hammer, J.A., Liu, Z., et al. (2014). Lattice light-sheet microscopy: imaging molecules to embryos at high spatiotemporal resolution. Science 346, 1257998. 10.1126/science.1257998.

35. Barbet, G., Nair-Gupta, P., Schotsaert, M., Yeung, S.T., Moretti, J., Seyffer, F., Metreveli, G., Gardner, T., Choi, A., Tortorella, D., et al. (2021). TAP dysfunction in dendritic cells enables noncanonical cross-presentation for T cell priming. Nat. Immunol. 22, 497–509. 10.1038/s41590-021-00903-7.

36. Freund, E.C., Lock, J.Y., Oh, J., Maculins, T., Delamarre, L., Bohlen, C.J., Haley, B., and Murthy, A. (2020). Efficient gene knockout in primary human and murine myeloid cells by non-viral delivery of CRISPR-Cas9. J. Exp. Med. 217, e20191692. 10.1084/jem.20191692.

37. Johannessen, L., Sundberg, T.B., O’Connell, D.J., Kolde, R., Berstler, J., Billings, K.J., Khor, B., Seashore-Ludlow, B., Fassl, A., Russell, C.N., et al. (2017). Small-molecule studies identify CDK8 as a regulator of IL-10 in myeloid cells. Nat. Chem. Biol. 13, 1102–1108. 10.1038/nchembio.2458.

38. Griffin, G.K., Wu, J., Iracheta-Vellve, A., Patti, J.C., Hsu, J., Davis, T., Dele-Oni, D., Du, P.P., Halawi, A.G., Ishizuka, J.J., et al. (2021). Epigenetic silencing by SETDB1 suppresses tumour intrinsic immunogenicity. Nature 595, 309–314. 10.1038/s41586-021-03520-4.

39. Jaber, M., Radwan, A., Loyfer, N., Abdeen, M., Sebban, S., Khatib, A., Yassen, H., Kolb, T., Zapatka, M., Makedonski, K., et al. (2022). Comparative parallel multi-omics analysis during the induction of pluripotent and trophectoderm states. Nat. Commun. 13, 3475. 10.1038/s41467-022-31131-8.

