## Supplemental Figures for "Submicron-Sampling of Living Cells by Macrophages"

**Submicron-Sampling of Living Cells by Macrophages – Supplementary Information**

**Contents**

**Supplementary Figures**

Supplementary Fig. 1: Uncropped gel related to Fig. 5
Supplementary Fig. 2: Tissue gating strategies related to Figure 1 and Extended Data Fig. 1
Supplementary Fig. 3: Lymph node, tumor, and spleen gating strategies related to Figure 1 and Extended Data Fig. 1
Supplementary Fig. 4: Gating strategy for antigen transfer assay and toxicity/efficacy for drugs in Fig. 2
Supplementary Fig. 5: Gating strategy for vesicle flow, related to Fig. 4

**Supplementary Movie Legends**

Supplementary Movie 1: NSPARC confocal live imaging captures trogocytosis-like sampling
Supplementary Movie 2: Lattice light-sheet imaging captures trogocytosis-like sampling

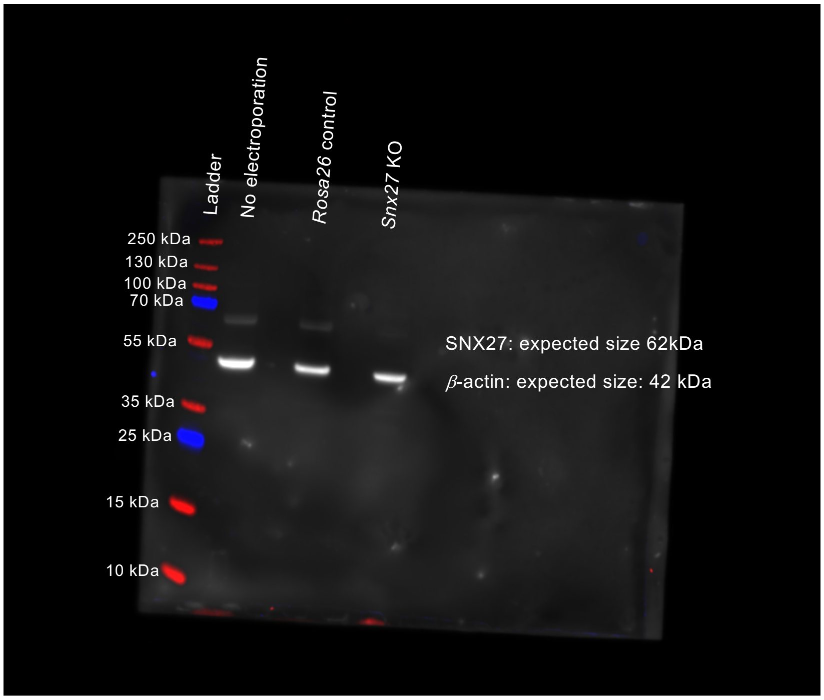


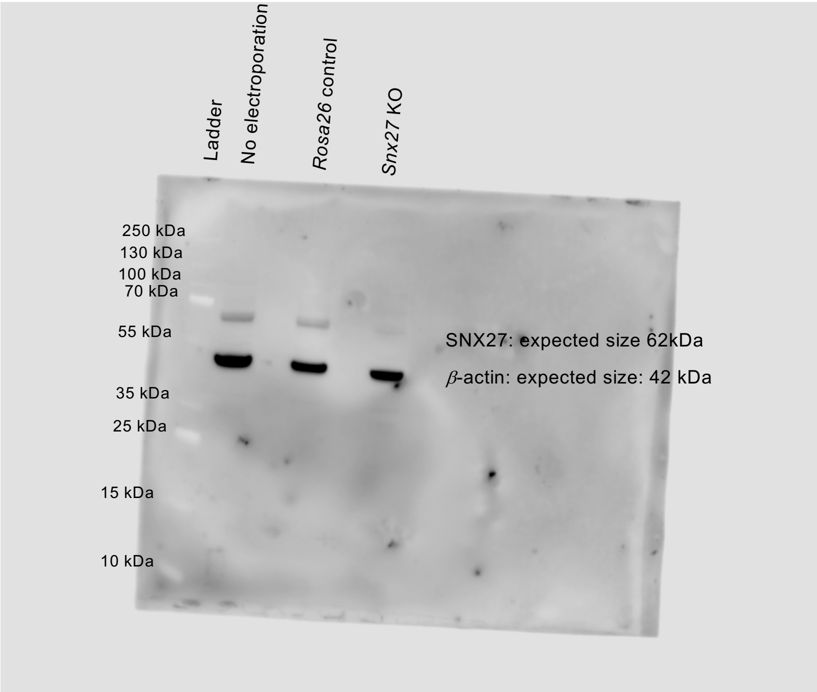


Controls were run on the same gel.

**Supplementary Fig. 1: Uncropped gel related to Fig. 5**

Protein lysate from unedited, *Rosa26*-targeted, and *Snx27* KO BMDMs were probed for SNX27 and β-actin.

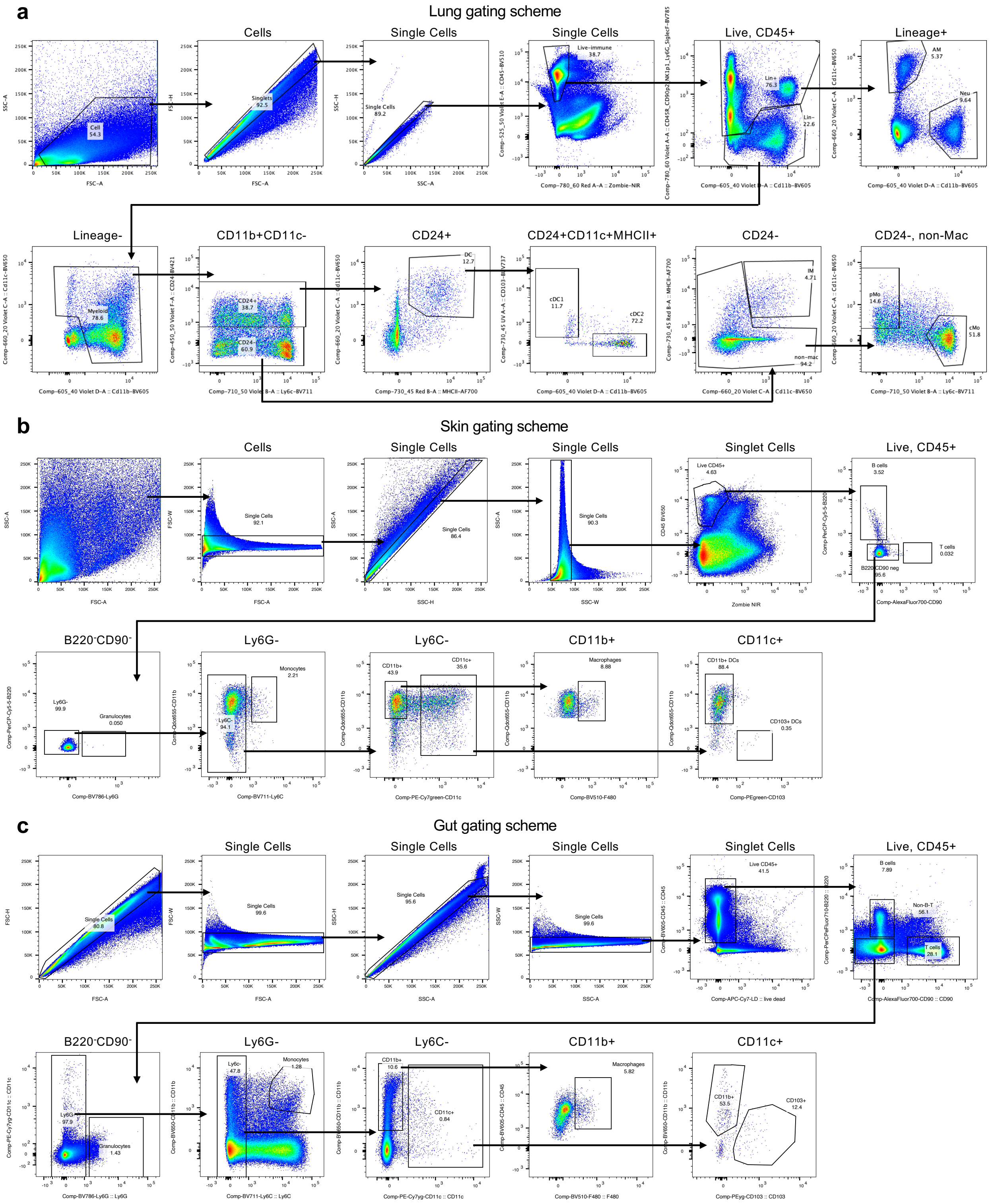


**Supplementary Fig. 2: Tissue gating strategies related to Figure 1 and Extended Data Fig. 1**
Representative flow cytometric gating strategy for myeloid populations in the lung (**a**), skin (**b**), and gut (**c**)

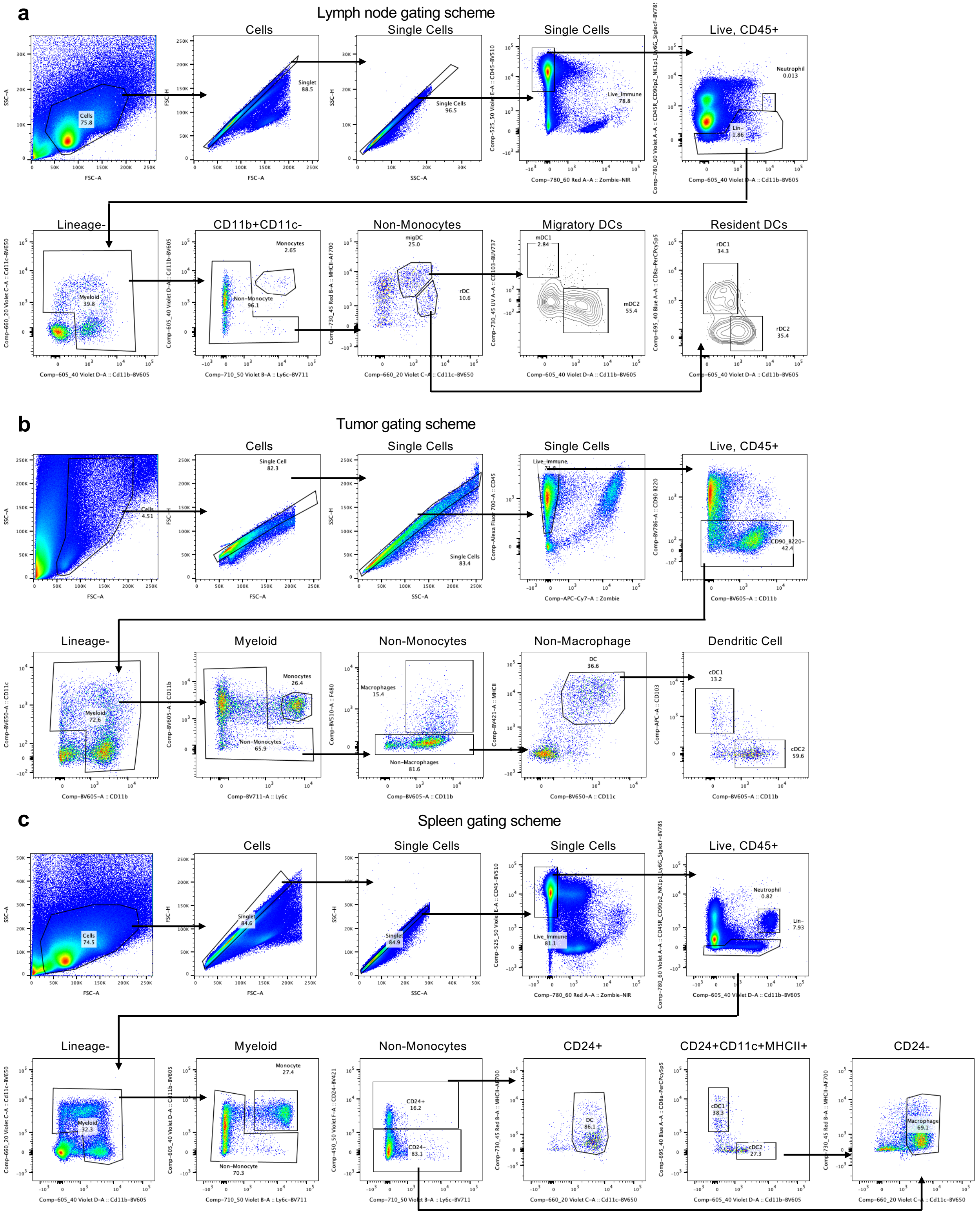


**Supplementary Fig. 3: Lymph node, tumor, and spleen gating strategies related to Figure 1 and Extended Data Fig. 1**

Representative flow cytometric gating strategy for myeloid populations in the lymph node (**a**), B16-ZsGreen tumors (**b**), and spleen (**c**)

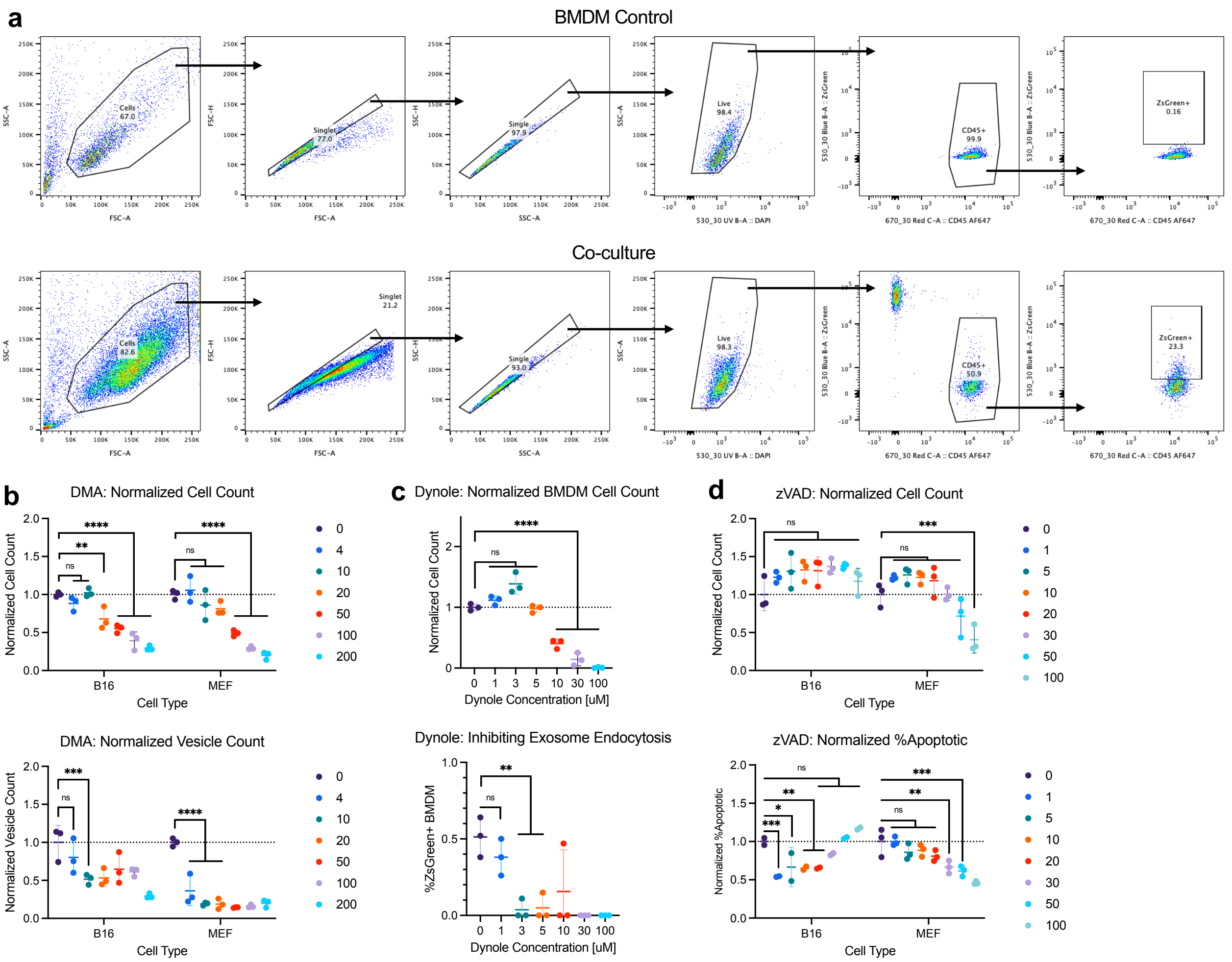


**Supplementary Fig. 4: Gating strategy for antigen transfer assay and toxicity/efficacy for drugs in Fig. 2**

**a,** Representative cytometric gating strategy for bone marrow derived macrophage (BMDM) co-culture with ZsGreen-expressing target cell and gating of ZsGreen positivity based on BMDM-alone control. **b,** B16-ZsGreen or MEF-ZsGreen target cells were cultured with DMA at indicated concentrations for 2 hours. Cell number was evaluated by flow cytometry to monitor toxicity (top) and supernatant was collected to quantify ZsGreen$^{+}$ vesicles by small particle flow cytometry. Teal arrows indicate concentration used in this study (10 uM DMA) and toxic concentrations are grayed out for functional studies. Shown are n = 3 technical replicates, mean +/- s.d. **c,** BMDMs were cultured with ZsGreen exosomes isolated using the ExoQuick kit with indicated concentrations of dynamin inhibitor Dynole 34-2. Cell number and exosome endocytosis were measured by flow cytometry using CountBright beads or quantifying %ZsGreen$^{+}$, respectively. Toxic concentrations are grayed out for functional studies. Shown are n = 3 technical replicates, mean +/- s.d. **d,** B16-ZsGreen or MEF-ZsGreen target cells were cultured with indicated concentrations of zVAD either in the absence (top) or presence (bottom) of Staurosporine to evaluate zVAD toxicity and efficacy, respectively. Cell number (top) and %AnnexinV$^{+}$DAPI$^{+/-}$ apoptotic cells (bottom) were evaluated by flow cytometry. Read arrows indicate concentration used in this study (20uM or 30 uM zVAD) and toxic concentrations are grayed out for functional studies. Shown are n = 3 technical replicates, mean +/- s.d.

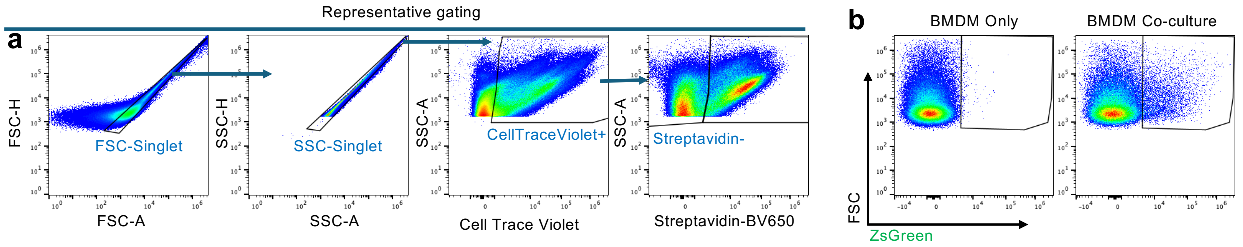


**Supplementary Fig. 5: Gating strategy for vesicle flow, related to Fig. 4**

**a,** Representative gating scheme to identify intracellular, protein-containing vesicles. **b,** Discrimination of ZsGreen+ vesicles allows for tracking of vesicles containing target-cell-derived protein.

### Supplementary Movie Legends

**Supplementary Movie 1: NSPARC confocal live imaging captures trogocytosis-like sampling**

Three-dimensional surface rendering of confocal live imaging of TdTomato+ BMDM and ZsGreen+ B16F10 interactions using the NSPARC detector visualizing the formation and separation of a vesicle from ZsGreen+ B16 into TdTomato+ BMDM. Time shown as m:ss. Z-stacks within series were captured in 30 sec intervals.

**Supplementary Movie 2: Lattice light-sheet imaging captures trogocytosis-like sampling**

Three-dimensional surface rendering of lattice light-sheet imaging of TdTomato+ BMDM and ZsGreen+ B16F10 capturing the stretching and separation of ZsGreen+ vesicle from target cell. Time shown as m:ss.
