## Extended Data for "Submicron-Sampling of Living Cells by Macrophages"

### Submicron-Sampling of Living Cells by Macrophages - Extended Data and Figure Legends

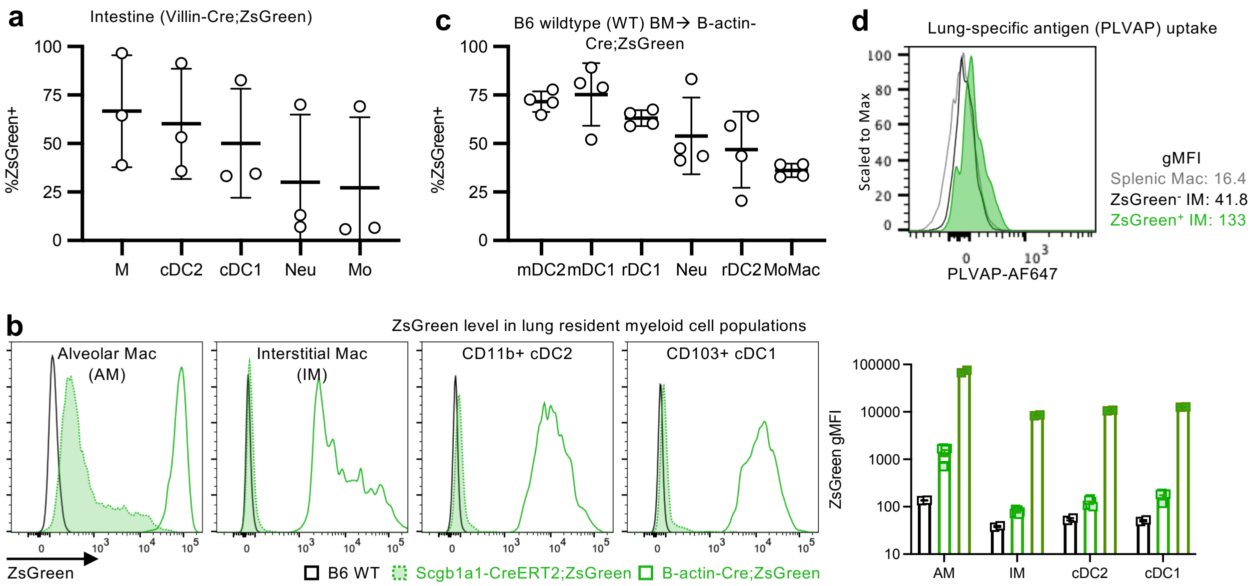


**Extended Data Figure 1: ZsGreen uptake from tissues is not due to Cre mis-expression.**
**a**, Gut taken from mice expressing ZsGreen under the Villin promoter. Isolation of myeloid cells from Villin-Cre;ZsGreen mice shows uptake in gut in macrophages as well as DCs. **b**, Representative flow (left) or quantification of ZsGreen (right) in lung myeloid cell populations in B6 wildtype mice, Scgb1a1-CreERT2;ZsGreen mice with lung-specific ZsGreen expression, and B-actin-Cre-ZsGreen mice. n = 2-6 mice per condition. **c**, Bone marrow transplant of B6 wildtype mice into B-actin-Cre;ZsGreen expressing mice. Myeloid cells isolated from skin-draining inguinal lymph nodes show ZsGreen sampling. **d**, Intracellular staining for airway specific PLVAP protein in lung (black) compared to splenic (gray) myeloid cell populations with ZsGreen+ lung macrophages (green) enriching for PLVAP signal. Shown are mean +/- standard deviation. Representative of 3 experiments, n = 3-6 mice per experiment. M, macrophage; Neu, neutrophil; Mo, monocyte; mDC2, migratory cDC2; mDC1, migratory cDC1; rDC1, residential cDC1; rDC2, residential cDC2; MoMac, Monocyte/Macrophage.

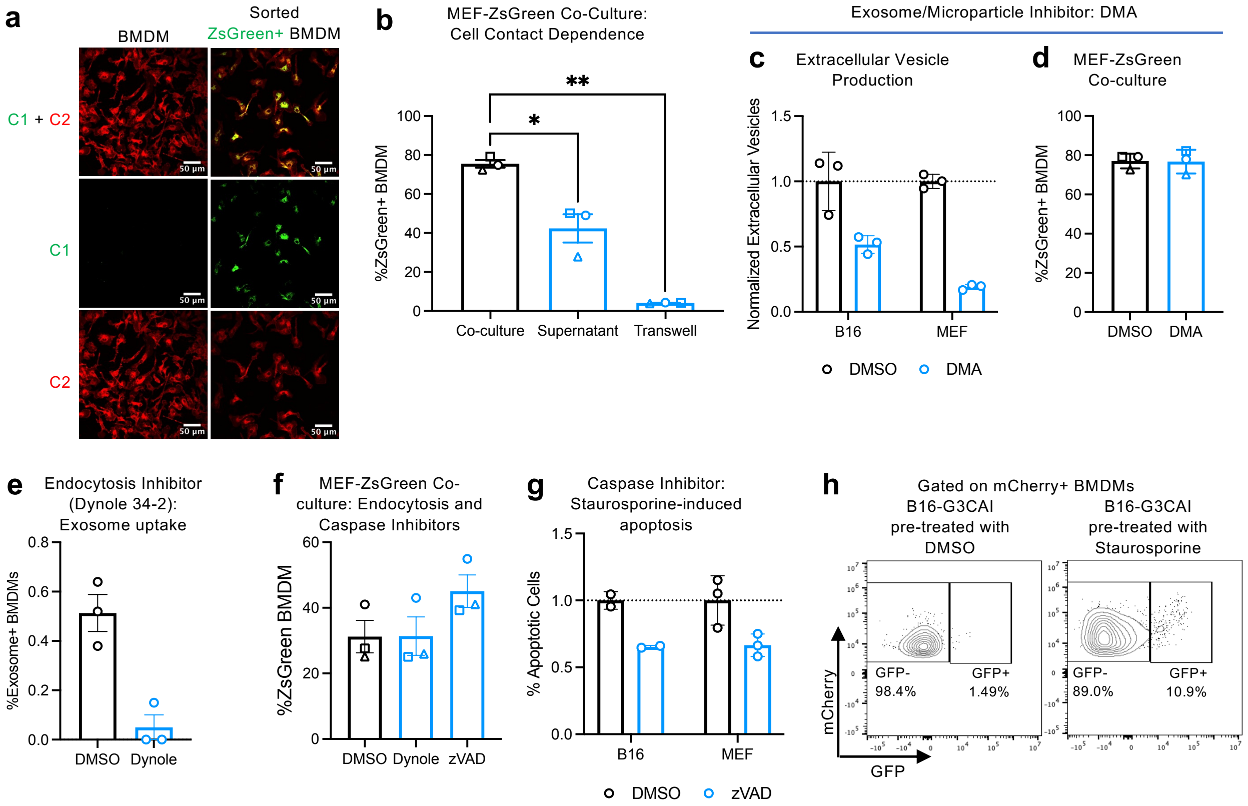


**Extended Data Figure 2: Live cells can be sampled in a cell-contact dependent manner without caspase activation**

**a,** Visualization of ZsGreen (C1) puncta ingested from target cells within membrane Tomato (C2) labeled BMDMs. ZsGreen+ BMDMs were sorted and imaged after co-culture with B16-ZsGreen cells. **b,** BMDMs were co-cultured directly with MEFs, with MEF supernatant, or with a transwell insert containing MEFs. **c-d,** Target cells (**c**) or BMDM-MEF co-cultures (**d**) were treated with DMA or DMSO vehicle control. Extracellular vesicle production in supernatant (**c**) or ZsGreen uptake (**d**) was measured by flow cytometry. **e,** Effect of Dynole on endocytosis of ZsGreen$^{+}$ exosomes by BMDMs. **f,** BMDM-MEF co-cultures were treated with indicated inhibitors or DMSO vehicle control. **g,** Target cells were treated with staurosporine and/or zVAD and AnnexinV$^{+}$DAPI$^{+/-}$ apoptotic cells were measured with flow cytometry. **h,** BMDM co-culture with B16-F10 expressing GFP caspase 3 activity indicator (B16-GC3AI) demonstrates uptake is predominantly from live cells. n = 3 biological replicates. Shown are mean of technical replicates +/- s.e.m. Paired t-test or one-way ANOVA. * $p < 0.05$, ** $p < 0.01$.

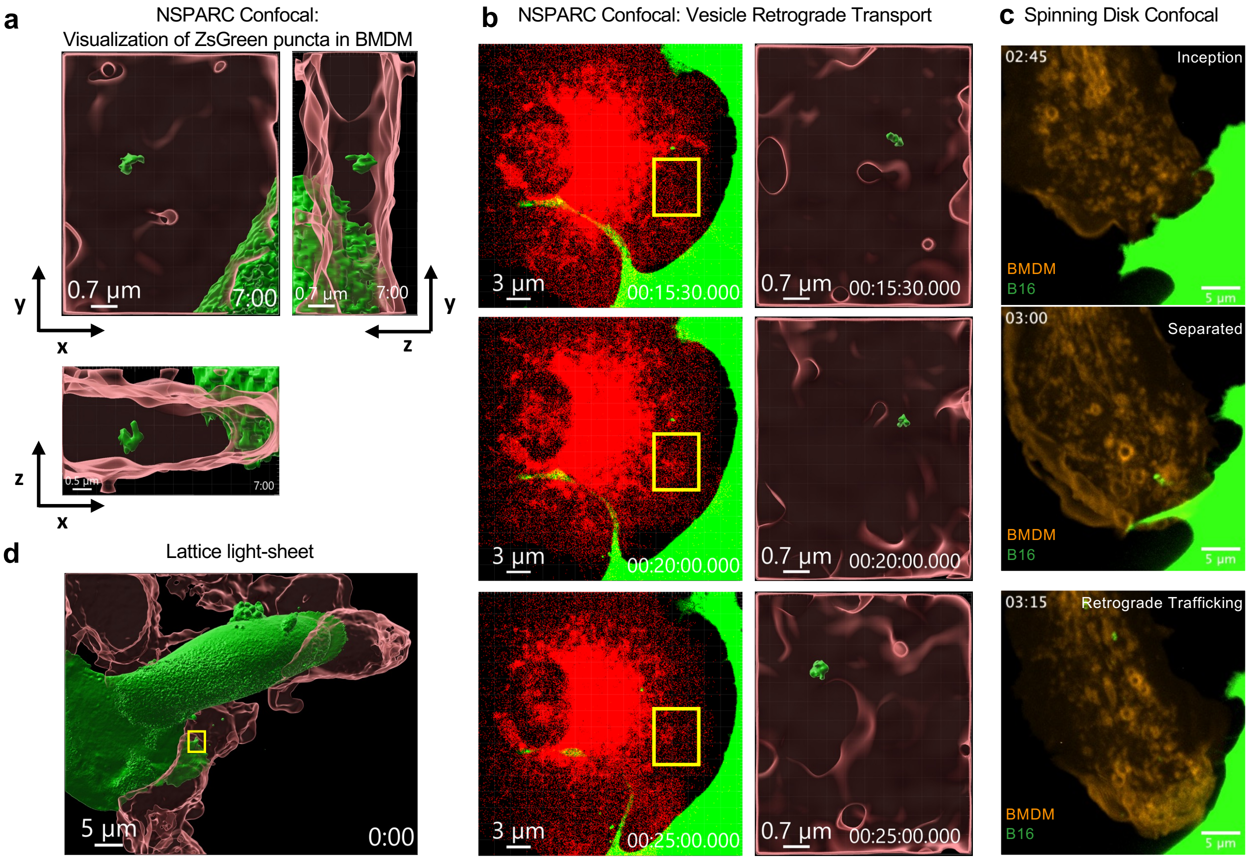


**Extended Data Figure 3: Nibbling can sample target cell protein**

**a,** Three-dimensional surface renderings from the YZ (left) and XY (bottom right) directions to identify intracellular localization of separated vesicle from same ROI as Fig. 3a, b. **b,** Two-color volume rendering showing interactions between TdTomato+ BMDM and ZsGreen+ B16(left) captured immediately after the series in Fig 3b. Three-dimensional surface renderings(right) of ROIs show separated vesicle trafficking towards the nucleus. Time shown as hh:mm:ss. **c,** Spinning-disk confocal imaging of ZsGreen+ B16 and TdTomato+ BMDM showing ingested ZsGreen+ puncta within BMDMs. Time shown as hh:mm. **d,** Three-dimensional surface rendering from Lattice light-sheet imaging of TdTomato+ BMDM and ZsGreen+ B16

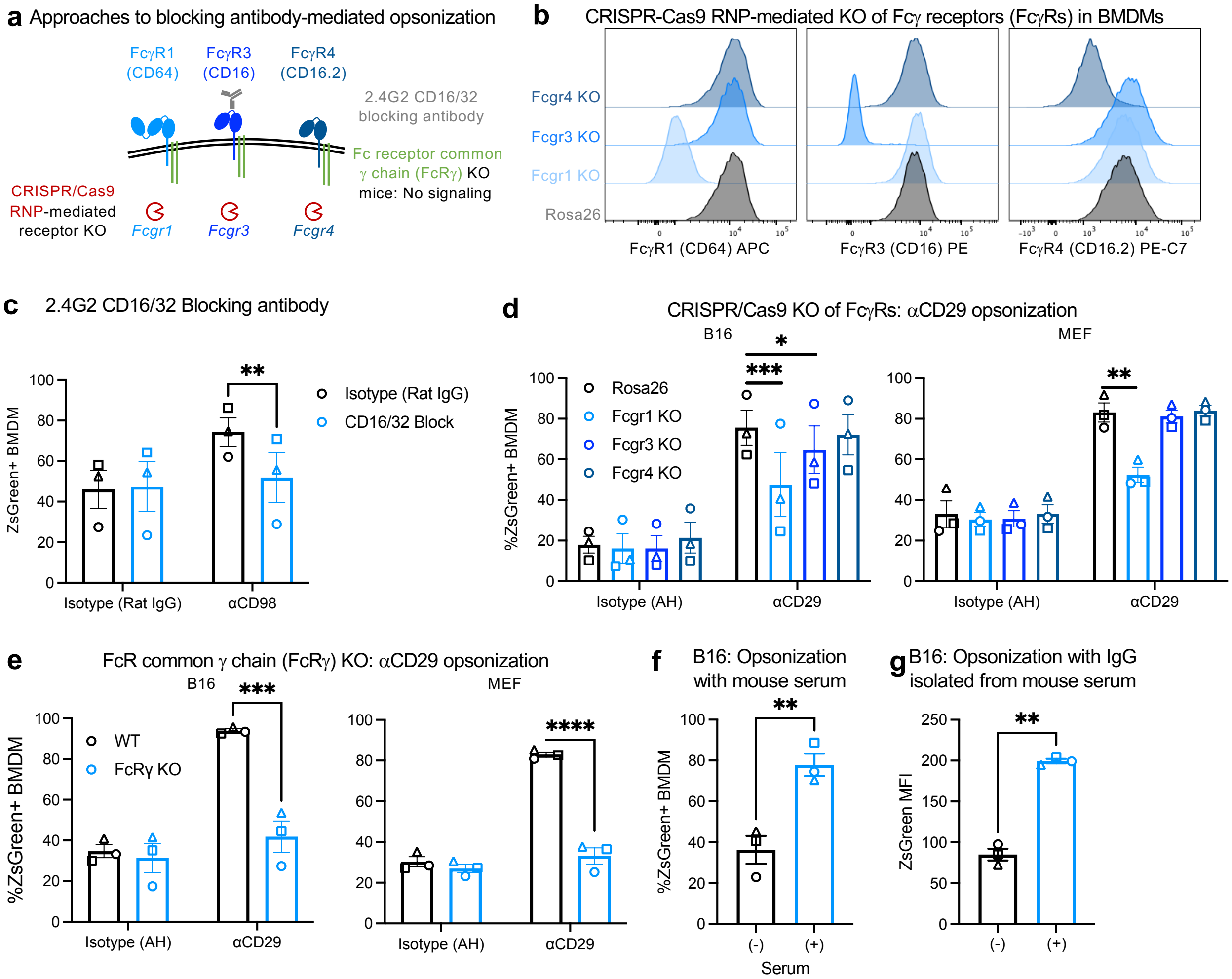


**Extended Data Figure 4: Antibody opsonization amplifies trogocytosis**

**a,** Schematic outlining methods used to block or knock-out (KO) FcgR binding and signaling. **b,** BMDMs targeted at the Rosa26 control locus or receptor loci using CRISPR-Cas9 RNP. **c,** BMDMs were pre-incubated with 10 ug/mL 2.4G2 CD16/32 blocking antibody or Rat IgG2b prior to co-culture with antibody-opsonized B16-ZsGreen target cells. **d-e,** CRISPR-targeted BMDMs (d) or BMDMs isolated from B6 wildtype or FcgR KO (e) mice were co-cultured with B16-ZsGreen or MEF-ZsGreen target cells pre-coated with antibody. **f-g,** BMDMs co-cultured with B16-ZsGreen pre-coated with serum (f), or IgG isolated from mouse serum (g). n = 3 biological replicates. Shown are mean of technical replicates +/- s.e.m. Paired t-test. * $p < 0.05$, ** $p < 0.01$.

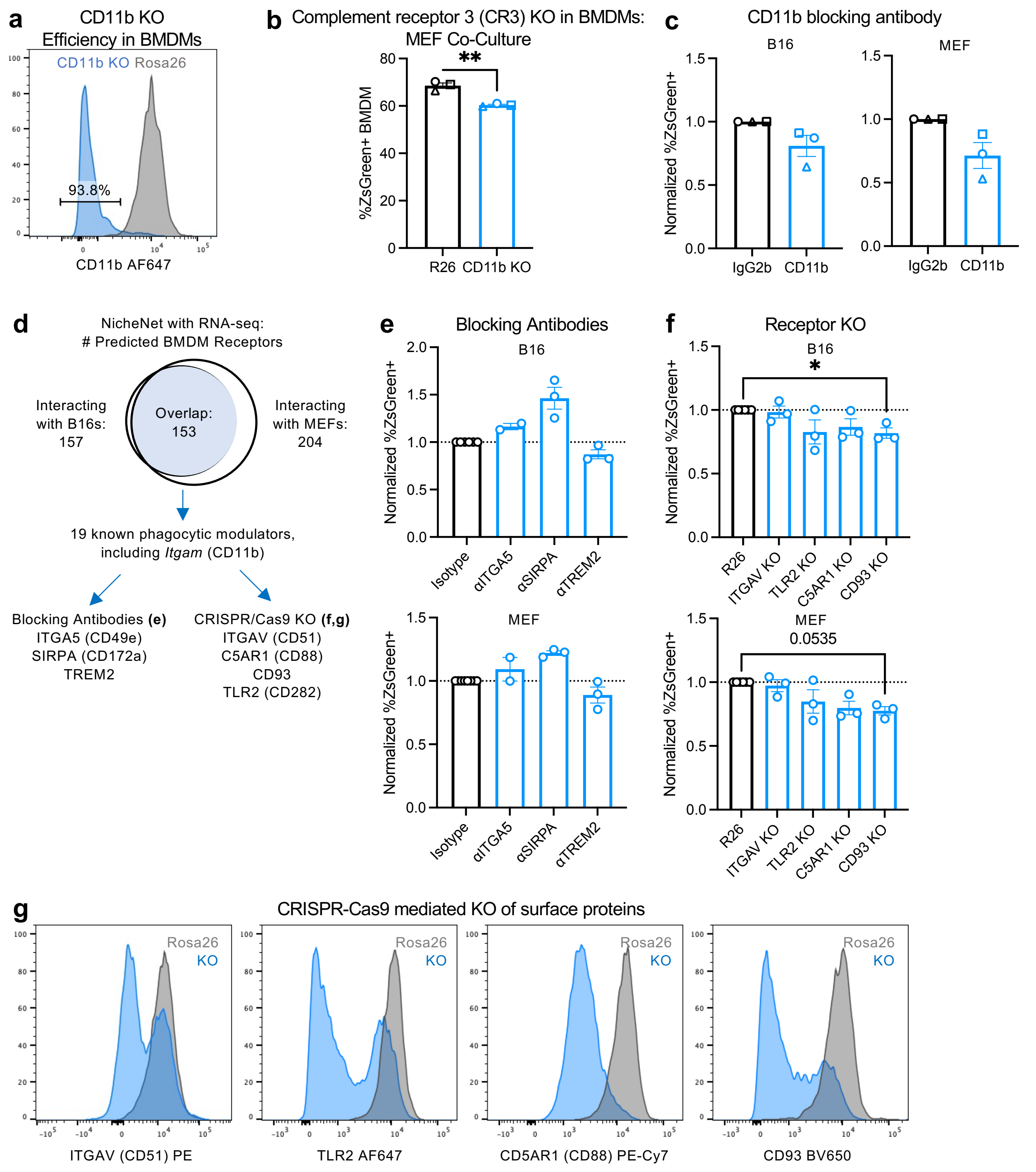


**Extended Data Figure 5: Knockout of CD11b and CD93 decreases live sampling from B16-ZsGreen target cells**

**a,** Representative protein knockdown 4 days after CRISPR/Cas9-mediated gene targeting. **b,** ZsGreen uptake in KO cells compared to Rosa26 (R26) controls. **c,** BMDMs were pre-treated with 1ug/uL IgG2b or CD11b-blocking antibody for 1 hour before 16-hour co-culture antigen transfer assays. **d,** Schematic outlining strategy to identify potential modulators of live sampling. **e,** BMDMs were co-cultured with B16-ZsGreen or MEF-ZsGreen target cells for 16 hours with 5 ug/mL blocking antibody or the appropriate isotype control prior to flow cytometric analysis of

ZsGreen uptake. **f-g** BMDMs targeted at the *Rosa26* control locus or receptor loci using CRISPR-Cas9 RNP and co-cultured with B16-ZsGreen or MEF-ZsGreen target cells for 16 hours prior to flow cytometric analysis of ZsGreen uptake (**f**) and receptor expression (**g**). Shown are mean +/- s.e.m. n = 2-3 experiments per receptor, 2-3 technical replicates per experiment. Kruskal-Wallis test with Dunn's multiple comparisons test. * $p < 0.05$.

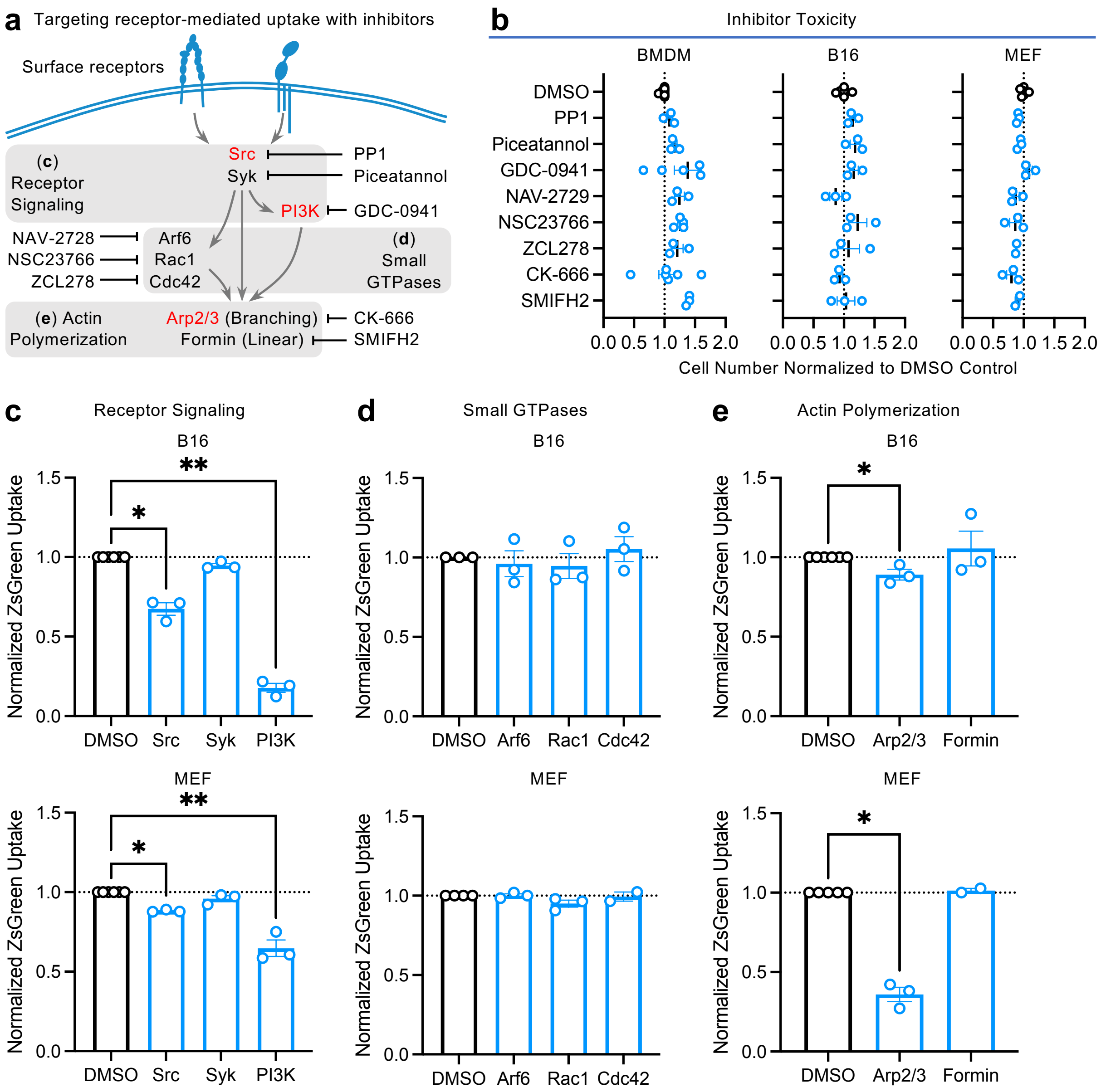


**Extended Data Fig. 6: Src, PI3K, and Arp2/3 inhibition decrease live sampling**

**a,** Schematic illustrating inhibitors used to target components of receptors-mediated uptake. **b-e,** BMDMs, B16-ZsGreen target cells, and MEF-ZsGreen target cells were cultured alone or co-culture for antigen transfer assay and treated with DMSO control, 10 uM PP1, 5 uM Piceatannol, 10 uM GDC-0941, 1.35 uM NAV-2729, 25 uM NSC23766, 10 uM ZCL278, 200 uM CK-666, or 5 uM SMIFH2 for 16 hours prior to flow cytometry analysis for cell number (**b**) or ZsGreen uptake (**c-e**). Data are normalized to DMSO control and compiled over 3-6 experiments. Shown are mean +/- s.e.m. Kruskal Wallis test and Dunn's multiple comparisons. *, $p < 0.05$; **, $p < 0.01$.

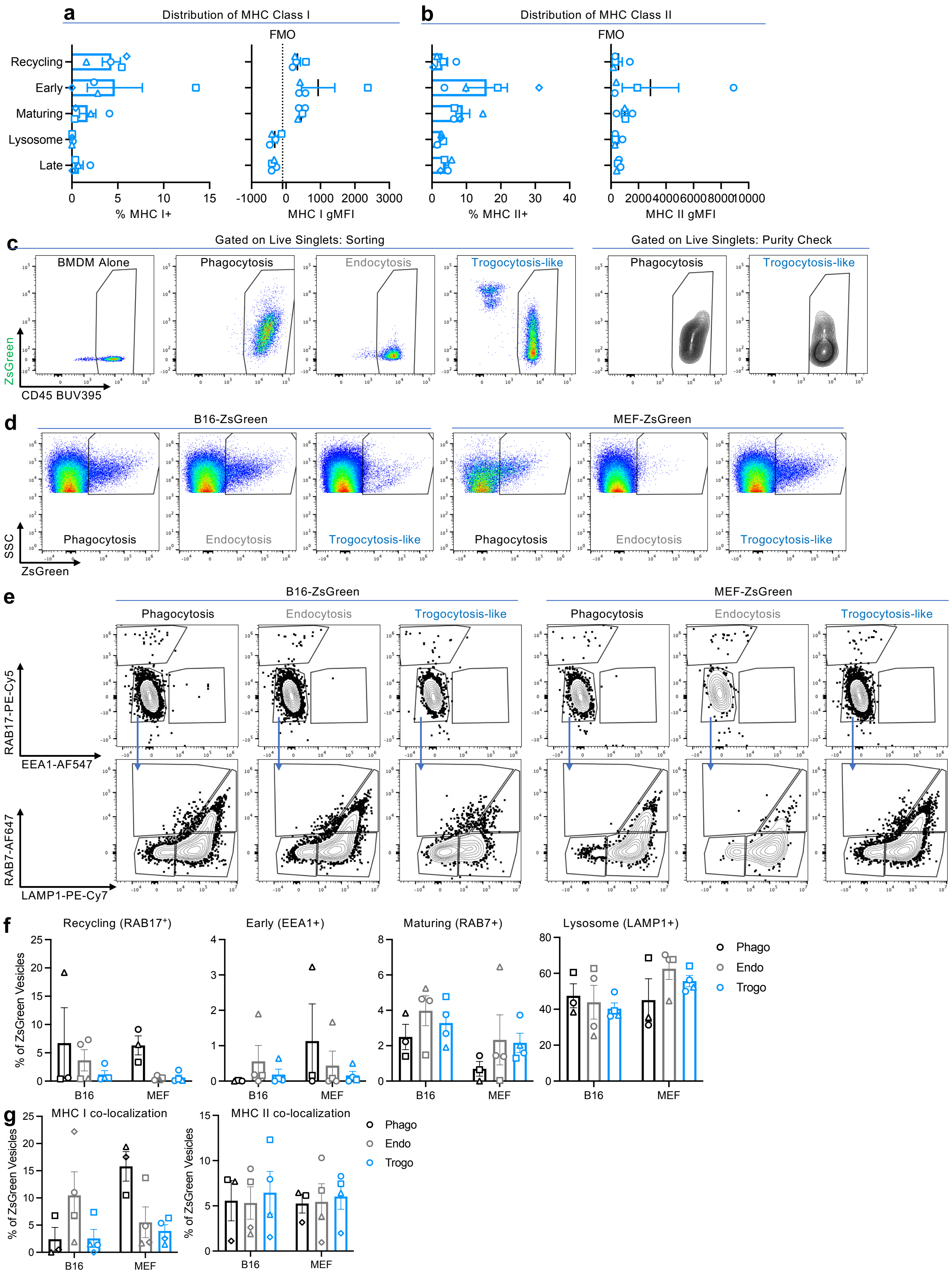


**Extended Data Figure 7: Live-cell associated antigen fills a discrete vesicular compartment**

**a-b,** Distribution of MHC I **(a)** and MHC II **(b)** in identified vesicular compartments inclusive of both ZsGreen+ and ZsGreen- vesicles. n = 3-4 biological replicates. Shown are mean of technical replicates +/- s.e.m. **c,** Representative sorting gate and purity check for CD45+ BMDMs from 16-hour co-culture. **d,** Representative ZsGreen+ gating from different vesicle flow preps gated on CellTraceViolet+Streptavidin- intracellular vesicles. **e,** Representative flow cytometry of vesicle populations from phagocytosis, endocytosis, and trogocytosis of B16-ZsGreen or MEF-ZsGreen target cells. **f,** Summary quantification of proportion of ZsGreen+ vesicles in different vesicular compartments. **g,** Proportion of ZsGreen+ vesicles that were either MHC I+ or MHC II+. n = 3-4 biological replicates. Shown are mean of technical replicates +/- s.e.m.

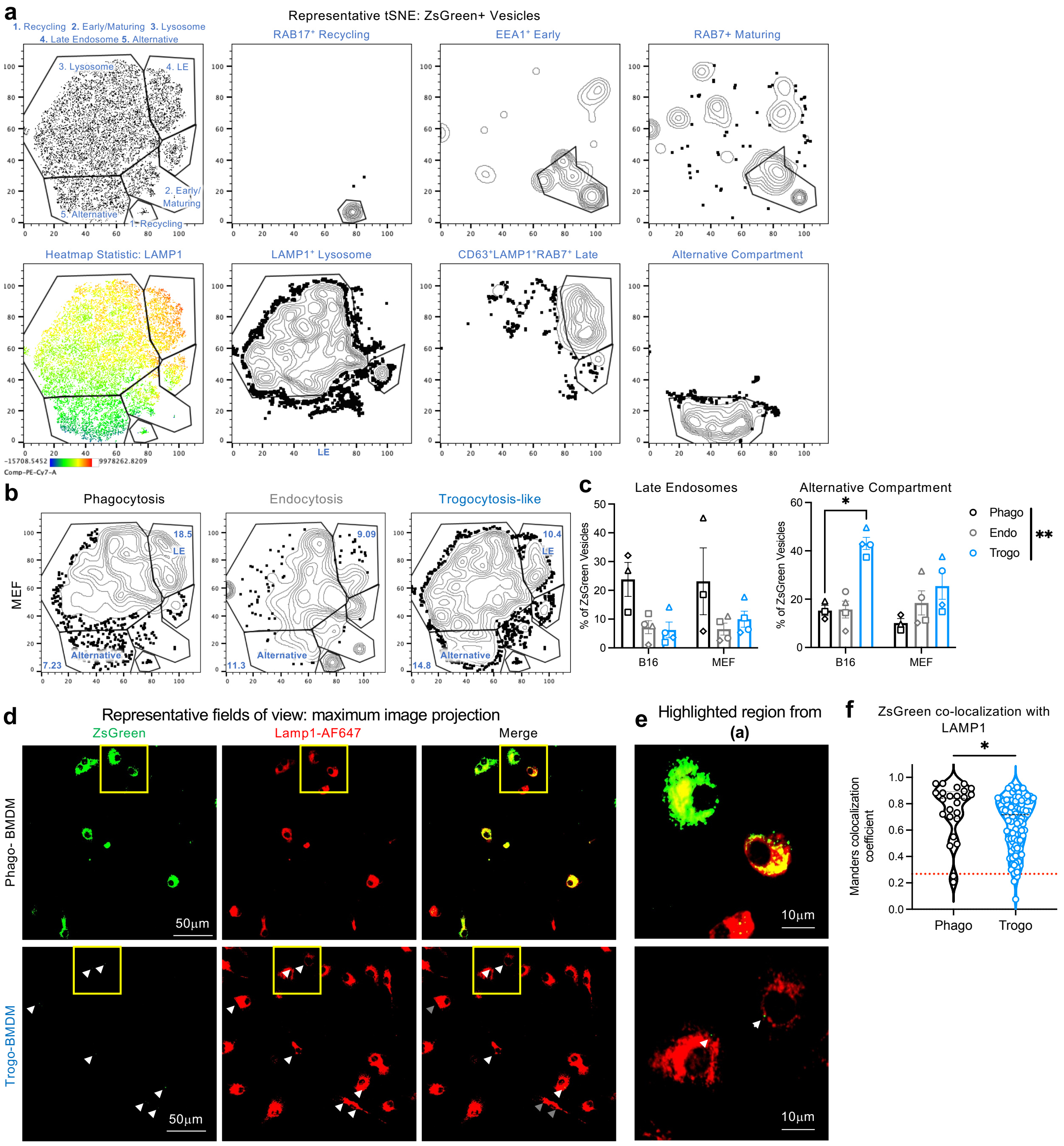


**Extended Data Figure 8: Live-cell associated antigen fills a LAMP1-negative vesicular compartment**

**a,** tSNE of concatenated ZsGreen+ vesicles from one experiment demonstrating gating of different vesicular compartments. **b,** Distribution of ZsGreen+ vesicles in derived from different sampling mechanism. **c,** Summary quantification of proportion of ZsGreen+ vesicles in late endosome and alternative vesicular compartment gates by gating on tSNE populations. n = 3-4 biological replicates. Shown are mean of technical replicates +/- s.e.m. **d-f,** Sorted ZsGreen+ BMDMs after co-culture with killed B16-ZsGreen (Phago-BMDM) and live B16-ZsGreen (Trogo-BMDM) were fix, permeabilized, and stained with LAMP1-AF647. Representative images are shown in (**d,e**) and quantification per field of view is shown in (**f**). Phago, n = 24; Trogo, n=96. Two-sided student's t-test or Two-way ANOVA. * $p < 0.05$, ** $p < 0.01$.

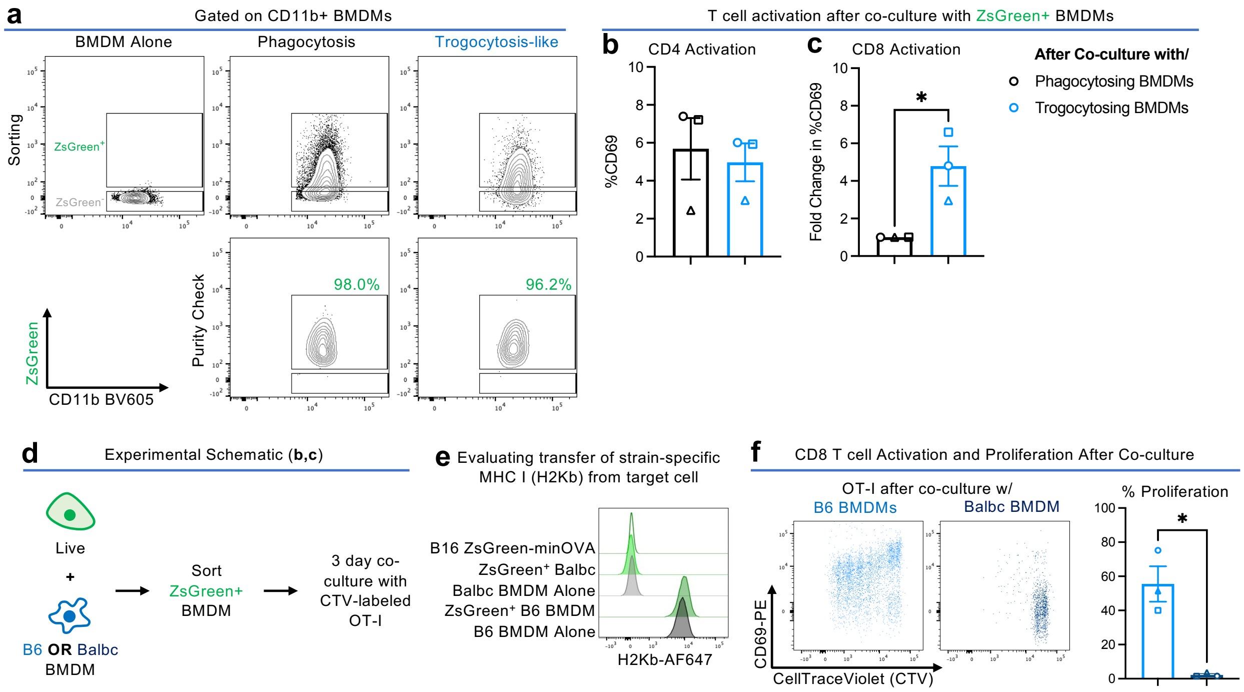


**Extended Data Figure 9: Macrophages process live-cell associated antigen to affect CD8 T cell activation**

**a,** Representative flow plots showing sorting and purity of ZsGreen+ BMDMs after co-culture with live or apoptotic B16F10-ZsGreen-minOVA target cells. **b-c,** Activation and proliferation of OT-I CD8 T cells (**b**) and OT-II CD4 T cells (**c**) as read out by CD69 expression after 3 days of co-culture with ZsGreen+ BMDMs from phagocytosis (black) or trogocytosis (blue) conditions. **d**, Experimental schematic to evaluate strain specific MHC-I transfer and T cell activation by ZsGreen$^{+}$ B6 or Balbc-derived BMDMs. **e**, H2-Kb expression on B16 ZsGreen-minOVA target cells; ZsGreen+ BMDMs after co-culture with B16 ZsGreen-minOVA cels, and BMDMs cultured alone. **f**, OT-I CD8 T cell proliferation after 3-day co-culture with FACS-isolated ZsGreen+ B6 or Balb/c BMDMs. n = 3 biological replicates with n = 3 technical replicates. Shown are mean of technical replicates +/- s.e.m. Two-sided paired t-test. * $p < 0.05$.

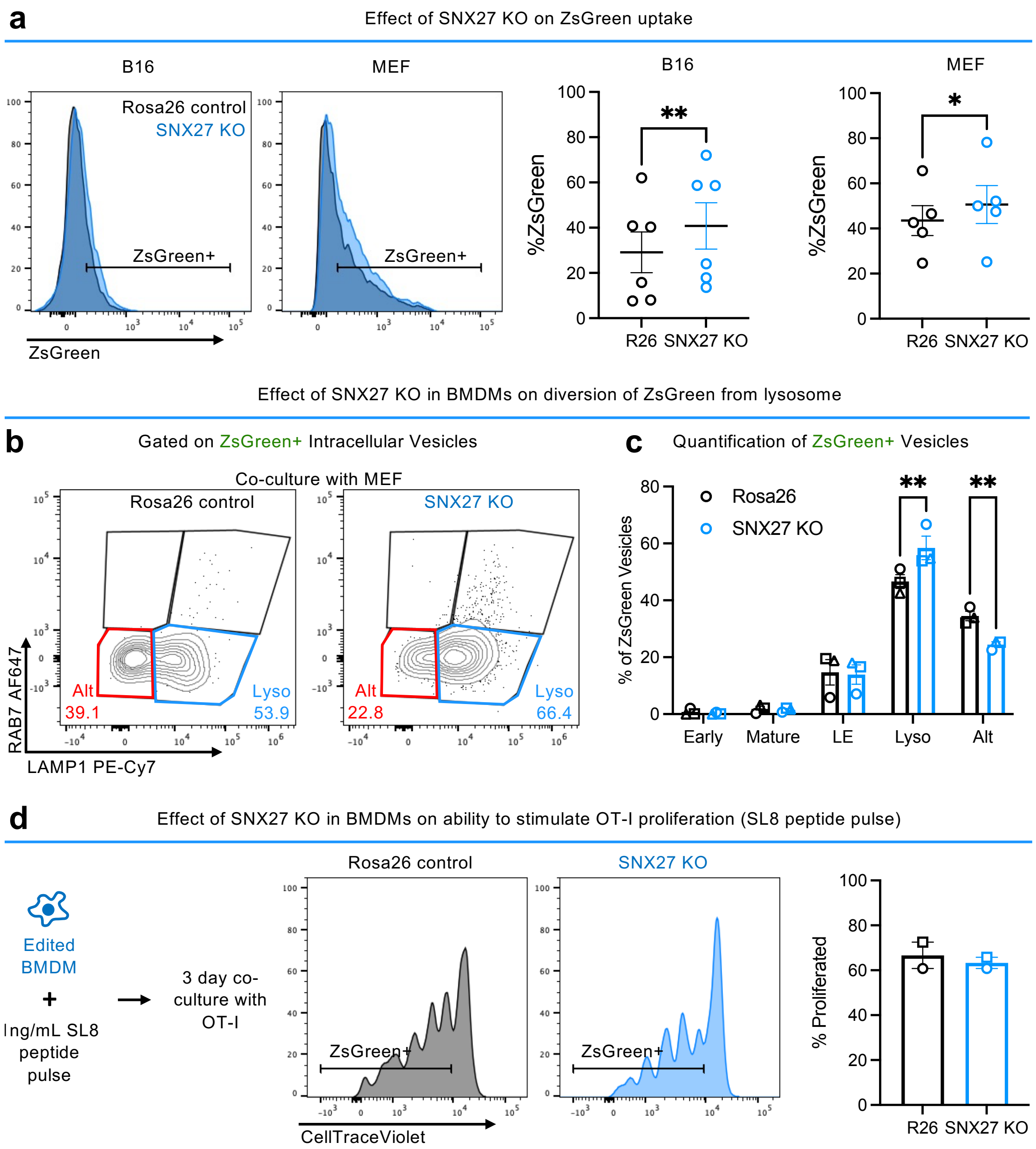


**Extended Data Fig. 10: SNX27 KO reduces antigen diversion from the lysosome but not uptake or presentation capacity**

**a-c,** ZsGreen uptake (**a**) and vesicle distribution (**b,c**) in SNX27 KO cells compared to Rosa26 (R26) controls. **d,** OT-I proliferation after 3-day co-culture with peptide-pulsed SNX27 KO and Rosa26 control BMDMs.
